# Phenotypic and genotypic consequences of CRISPR/Cas9 editing of the replication origins in the rDNA of *Saccharomyces cerevisiae*

**DOI:** 10.1101/647495

**Authors:** Joseph C. Sanchez, Anja Ollodart, Christopher R. L. Large, Courtnee Clough, Gina M. Alvino, Mitsuhiro Tsuchiya, Matthew Crane, Elizabeth X. Kwan, Matt Kaeberlein, Maitreya J. Dunham, M. K. Raghuraman, Bonita J. Brewer

## Abstract

The complex structure and repetitive nature of eukaryotic ribosomal DNA (rDNA) is a challenge for genome assembly, and thus, the consequences of sequence variation in rDNA remain unexplored. However, renewed interest in the role that rDNA variation may play in diverse cellular functions, aside from ribosome production, highlights the need for a method that would permit genetic manipulation of the rDNA. Here, we describe a CRISPR/Cas9 based strategy to edit the rDNA locus in the budding yeast *Saccharomyces cerevisiae.* Using this approach, we modified the endogenous rDNA origin of replication in each repeat by deleting or replacing its consensus sequence. We characterized the transformants that have successfully modified their rDNA locus and propose a mechanism for how CRISPR/Cas9 mediated editing of the rDNA occurs. In addition, we carried out extended growth and life span experiments to investigate the long-term consequences that altering the rDNA origin of replication has on cellular health. We find that long-term growth of the edited clones results in faster growing suppressors that have acquired segmental aneusomy of the rDNA containing region of chr XII or aneuploidy of chromosomes XII, II, or IV. Furthermore, we find that all edited isolates suffer a reduced life span, irrespective of their levels of extrachromosomal rDNA circles. Our work demonstrates that it is possible to quickly, efficiently and homogeneously edit the rDNA locus via CRISPR/Cas9. It serves as a model for modifying other parts of the rDNA and, more generally, for editing other tandemly repeated sequences in higher eukaryotes.

## Introduction

While the role of ribosomal DNA (rDNA) in ribosome production is uncontested, there has been renewed interest in exploring the role that rDNA variation may play in cell cycle regulation, lifespan and cancer (Wang and Lemos 2017; Parks *et al*. 2018). Because ribosomal content is one of the dominant components of cellular biomass, ribosomal RNA (rRNA) transcription imposes significant constraints on the speed with which cells can divide. The heavy transcriptional burden placed on the rDNA has resulted in a necessity for large numbers of repeated units to cope with the demand for rRNAs. This burden, and the inherently repetitive nature of these sequences can result in variation within an otherwise isogenic population. However, variation in the sequence or copy number of rDNA repeats has been difficult to assess experimentally (McStay 2016). Even though whole genome sequencing (WGS) data are available for hundreds to thousands of different eukaryotes, the large size of the repeated, homogeneous, tandem structure makes identifying and testing sequence and copy number variants challenging in most species. The copy number of repeats can be highly variable from individual to individual (Stults *et al*. 2008; Xu *et al*. 2017) and is often difficult to reliably determine when orthogonal methods, such as ddPCR or qPCR, quantitative hybridization and WGS read depth measurements give variable estimations of copy number from the same DNA sample (Xu *et al*. 2017; Chestkov *et al*. 2018). Adding to the complexity of the problem is the fact that in many eukaryotes, the rDNA is also found on multiple chromosomes. Directed mutational analysis, a technique so powerful for analyzing single copy sequences, has not been a realistic option in the rDNA of most organisms because of the difficulty in mutagenizing all the repeats simultaneously. Therefore the systematic testing of individual sequence or copy number variants has been genetically intractable.

In the budding yeast *Saccharomyces cerevisiae* limited mutational analysis of the rDNA locus has been carried out by taking advantage of a recessive hygromycin resistant mutation (T to C) at position 1756 of the 18S rRNA. When this variant rDNA repeat is introduced into yeast on a multi-copy plasmid, cells can become hygromycin resistant by deleting most or all of the chromosome XII copies of the rDNA (Chernoff *et al*. 1994; Wai *et al*. 2000) and relying on the plasmid copies of the rDNA for ribosome production. Using a plasmid shuffle protocol or direct reintegration of sequences into chr XII, researchers can examine the consequences of mutated rDNA elements. This basic protocol has been used to explore transcriptional regulatory elements of the PolI promoter and enhancer (Wai *et al*. 2000), the role of expansion segments in ribosome fidelity (Fujii *et al*. 2018), the sequence requirements for replication fork blocking at the 3’end of the 35S transcription unit (the replication fork barrier, RFB; Ganley *et al*. 2009; Saka *et al*. 2013), and on a more limited scale, the requirements for the rDNA origin of replication in copy number expansion (Ganley *et al*. 2009). While these experiments are elegant, there are problems with interpreting resultant phenotypes—in particular the introduction of altered rDNA sequences requires the simultaneous introduction of a selectable marker that becomes part of each of the amplified variant rDNA repeats.

Here, we edited the yeast rDNA locus directly without the introduction of a selectable marker by using CRISPR/Cas9 (Doudna and Charpentier 2014). We altered the origin of replication that is found in each repeat by deleting it entirely or by replacing the 11 bp A+T-rich origin consensus sequence with a G+C block or other functional origins. We describe our protocol for editing the rDNA, characterize the surviving clones and propose a mechanism by which transformants are able to successfully replace multiple tandem repeats. Among the surviving transformants complete sequence replacements are surprisingly frequent and easily detected. In addition, we analyzed the phenotypic consequence of altering the sequence of the rDNA origin. While our work was underway, a preliminary report from Chiou and Armaleo indicated that they had successfully inserted a 57 bp intron into the 18S rRNA gene in yeast using CRISPR/Cas9 technology (Chiou and Armaleo 2018), but they provided little phenotypic or genetic analysis of their transformants.

We find that deletion of the rDNA origin limits the number of rDNA repeats that can be maintained by passive replication to ~10. The resulting limitation on ribosome production creates a strong selective pressure for suppressors of the slow growth phenotype. Using multiple, long-term growth regimes, we find that suppressors arise through segmental aneusomy of the region of chr XII that contains the rDNA locus or through aneuploidy of chromosomes XII, II or IV. Replacing the rDNA origin with other functional origins or re-introducing the rDNA origin also leads initially to reduced rDNA copy number. Over ~140 generations, the locus re-expands but does not reach the initial ~150 copies. All strains that survived CRISPR editing at the rDNA locus showed a reduction in lifespan that is unrelated to the level of extrachromosomal rDNA circles (ERCs), which we found can vary among the strains by ~100-fold. Our genetic analysis of this understudied, essential, complex locus in yeast adds critical understanding to consequences of rDNA origin variation and serves as a proof of principle for editing the rDNA in other eukaryotes.

## Materials and Methods

### Strains, growth medium, plasmids, and PCR

All edited strains were derived from BY4741 or BY4742. Strains were verified by PCR, restriction digestion and/or Sanger sequencing of the PCR products.

Cells were grown at 30°C in defined synthetic complete medium (WFC), with or without uracil (for selection of the Cas9 plasmids) and supplemented with 2% dextrose.

pML104 (pML104; Laughery *et al*. 2015) was purchased from Addgene and cleaved sequentially with BclI and SwaI. Complementary oligos for each of the guide sequences were annealed and ligated into the cut pML104. Colonies were selected on LB-Amp plates and transformants were Sanger sequenced. Multiple guide plasmids were Gibson-assembled (Gibson 2011) from PCR products of single guide plasmids.

### Creating and characterizing the CRISPR/Cas9 edited rDNA arrays

To maximize the cutting potential of the system we cloned multiple guides into a single plasmid (pML104; Laughery *et al*. 2015) that contains the Cas9 gene, a *URA3* selectable marker, and the 2-micron origin of replication that stably maintains the plasmid at high copy number (pACD; Figure S1). We designed a 250 bp “gBlock” repair template with mutations at four possible NGG PAM sequences (rendering the repair template resistant to Cas9 cleavage) and a G+C block that replaced the 11 bp A+T-rich ARS consensus sequence (ACS; Figure 1B and Figure S1B). We transformed strain BY4741 with pML104 or pACD and included excess repair template in either single-stranded or double-stranded form. While pML104 produced hundreds of Ura+ colonies per microgram of plasmid DNA within three days, transformation with pACD with or without repair template produced only dozens of colonies of variable sizes that appeared over the course of a week’s incubation (Figure S2A). The reduced transformation efficiency is consistent with the assumption that unrepaired double stranded breaks in the rDNA are lethal.

**Figure 1:**
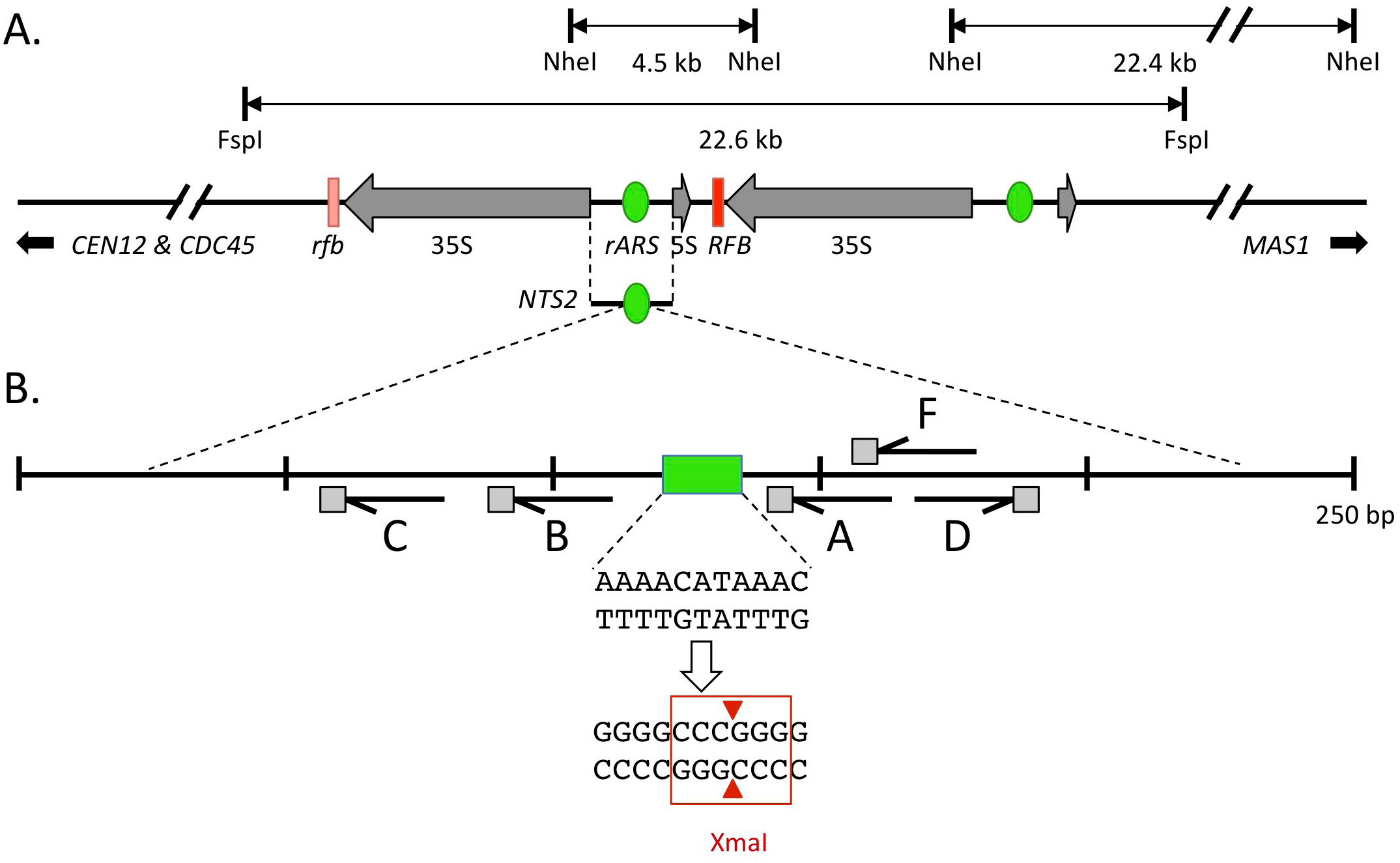
Maps of the rDNA locus and of features relevant for CRISPR/Cas9 editing. A) A map of the rDNA locus adapted from the *Saccharomyces* Genome Database (https://www.yeastgenome.org) representing just two of the ~150 rDNA repeats in the reference yeast strain. NheI and FspI restriction sites relevant to this study are indicated, as well as the proximal (*CEN12* and *CDC45*) and distal (*MAS1*) sequences used for hybridization probes. (The left RFB, indicated in light red, is a truncated version of the RFB that is present in all of the other repeats.) B) A 250 bp region of *NTS2* that contains the origin of replication. The A+T sequence of the *rARS* ACS (green box) was targeted for replacement by the G+C sequence that introduces an XmaI site. The half arrows indicate the guide RNA sequences and the gray squares indicate the NGG PAM sites.

After restreaking the transformants on selective plates (Figure S2B), we characterized their rDNA loci by performing PCR, restriction digestion (Figure S2C) and/or sequencing (Figure S3). Transformation with the repair template along with pACD was expected to produce no change in size of the PCR fragment if the target was edited correctly; however, the PCR product should contain an XmaI site that was absent in the original rDNA sequence (Figure 1B and Figure S1B). Four of the eight transformants that received pACD alone showed no signs of editing and the other four clones yielded reduced or no PCR fragments, suggesting they had suffered a deletion in the region (Figure S2C). Among a set of sixteen transformants that received both pACD and the repair template, we found a single clone with the appropriate PCR fragment size and sensitivity to XmaI as expected for correct editing (transformant #6; pACD + ss template; Figure S2C). The sequence of the PCR fragment revealed all four PAM mutations and the 11 bp G+C cassette (Figure S3A; refer to gBlock sequence in Table S1). All of the remaining transformants failed to produce a PCR fragment or produced a shorter than expected PCR fragment, suggesting that they had suffered a deletion. We sequenced the PCR fragment from one of these clones (transformant #2; pACD + ds template; Figure S2C) and found it to have a deletion that removed 167 bp of the origin region including all guide RNA sites (Table S1; Figure S3B). We repeated the transformation experiments, focusing primarily on small transformant colonies, and obtained and characterized in detail an additional *rARS^GC^* mutants (rARS^GC^−2) and an additional *rARS^Δ^* mutant (rARS^Δ^−2).

Editing of the rDNA repeats changed two features of the rDNA locus: the origin was removed and the copy number of the repeats was drastically decreased. To disentangle the phenotypes, we repeated the editing experiments using pACD and a template with the wild type rDNA origin and the four PAM site mutations. To detect introduction of *rARS^“wt”^* we took advantage of the fact that the sequence at two of the PAM sites could be distinguished by differential restriction enzyme digestion: an ApoI site is present at the edited PAM site associated with guide A and an HaeIII site is present at the native PAM site associated with guide D (Figure S1B; Figure S4A). PCR and restriction digest analysis revealed that three of the clones (lanes 2, 3 and 12) had a successful replacement with the modified “wt” rARS. We refer to these strains as rARS^“wt”^−2, rARS^“wt”^−3 and rARS^“wt”^−12.

To target *ARS1* or *ARS1^max^* to the rDNA locus, we included ~170 bp of *NTS2* homology flanking *ARS1* or *ARS1^max^* (Table S1). Using rARS^GC^−1 as the starting strain, we co-transformed *ARS1^max^* and the single guide RNA plasmid pF (Figure S1). We expected successful repair using the *rARS1^max^* template to produce a larger PCR fragment that was missing the XmaI site but instead would contain a BspHI site (Figure S1B). More than 50% of the slow-growing clones contained the desired ARS replacements (Figure S4B). We were also able to recover, at high efficiency, the *ARS1* or *ARS1^max^* replacements using plasmid pACD and BY4741 as the starting strain (Figure S4C). *ARS1* replacements produce the same larger sized PCR fragment that contains a novel BglII site (Figure S1B and Figure S4C).

### CHEF and 2D gel electrophoresis

For CHEF gel analysis of chromosome sizes, 1-1.5 ml of saturated yeast cultures were embedded in 0.5% GTG agarose plugs and processed for CHEF gel electrophoresis (Kwan *et al*. 2016). We used three CHEF gel running conditions in Bio Rad’s CHEF DRII: the full chromosome run (which separates all chromosomes except for IV and XII) was 64 hr in 0.8% LE agarose with 0.5X TBE at 165 volts with switch times from 47” to 170”; the rDNA run (which separates larger chromosomes, such as IV and XII) was 68 hr in 0.8% LE agarose with 0.5X TBE at 100 volts with switch times from 300” to 900”; the Fsp1 digested chromosomes were run as for the full chromosome run but for only 44 hours. To collect DNA for 2D gels, cells were grown to mid log phase, quick chilled in sodium azide and EDTA, and concentrated by centrifugation. Agarose plugs from frozen cell pellets were prepared by the protocol available at http://fangman-brewer.gs.washington.edu. Methods for restriction digestion of DNA in plugs and conditions for 2D gel electrophoresis of rDNA replication intermediates can be found in Sanchez et al. (2017). Southern transfer and hybridization with ^32^P-dATP PCR amplified probes were performed according to standard protocols. The split Southern/northern gel was performed as described in Sanchez, et al. (2017).

### Turbidostat growth

Turbidostats were constructed and run under software from (McGeachy *et al*. 2019). Samples were harvested at indicated times by collecting the effluent on ice over a ~15 minute interval. Custom R scripts were used to convert the calculated influent amount to a doubling time.

Run 1 parameters: Four separate colonies, two each of rARS^GC^−1 (colonies a and b) and rARS^Δ^−1 (colonies a and b), were grown in WFC medium. Each 200ml turbidostat was inoculated with 2 ml of an overnight culture. The cells were then allowed to grow until they reached ~OD_600_ = 1. At this point another sample was taken to manually measure the OD_600_ of the cells, which then was used to calibrate the turbidostat readings. The turbidostat was then set to maintain a constant OD_600_ of 1. At the same time each day, 15 ml of effluent from each turbidostat was collected on ice (approximately 10-15 minutes). Cells were pelleted at 4000 x g for 5 minutes and frozen at −20°C for downstream analysis. Glycerol stocks of 1 ml were also collected and stored at −80°C.

Run 2 parameters: The OD_600_ was set to 0.5 to more accurately reflect log phase growth, and 50 ml samples of cells were collected to account for the decreased density and to provide adequate sample for additional assays.

Doubling Time Analysis: The system measures the on/off cycling of the media pump that is delivering medium at a known rate. A custom R script fits a line to the change in the overall time the pump is on over a 2-hour window. The slope of that line is then scaled by the known flow rate and volume of the vessel. This calculation produces a doubling time for that 2 hour interval.

### Replicative Lifespan Analysis

Yeast replicative lifespans were performed as previously described (Kaeberlein *et al*. 2004). Briefly, strains were streaked from frozen stocks onto YEP glycerol media plates to minimize the chance of copy number changes in the rDNA. Strains were then patched onto YEP dextrose plates where they were allowed to grow for several generations. Virgin daughters were isolated, allowed to grow into mothers, and then individual daughters were manually removed. Each division of the mother cell was recorded until the mother cells failed to divide. Statistical significance was determined using Wilcoxon Rank-Sum analysis.

### Whole Genome Sequencing

Total cellular DNAs from rARS^GC^−1a and −1b samples were isolated from stationary yeast cells by standard methodology (Hoffman and Winston 1987). DNA was fragmented by Covaris sonication (450 peak factor, 30% duty factor, 200 cycles per burst) prior to single-index library construction using the KAPA Hyper Prep Kit. Following qPCR for quality control and quantification, libraries were sequenced on an Illumina NextSeq 500 using the manufacturer’s recommended protocols.

Libraries from the turbidostat cultures were prepared by isolating genomic DNA using the Hoffman-Winston protocol (Hoffman and Winston 1987). DNA quantification was assessed using Qbit. 50 ng aliquots of genomic DNA were prepared with the Nextera Library kit, according to kit instructions.

### Alignment and SNP/InDel Variant Calling

WGS reads were aligned using Bowtie2 (Bowtie/2.2.3) (Langmead and Salzberg 2012) or BWA-mem (BWA/ 0.7.15) (Li 2013) to the sacCer3 reference genome (Engel *et al*. 2014) depending on read length (BWA for >100bp and Bowtie2 for <100bp), then sorted and indexed using SAMtools/1.9 (Li *et al*. 2009). Duplicates were marked and removed using Picard tools (picard/2.6.0), resorted and indexed using SAMtools. The InDels in the alignments were realigned using the GATK/3.7 package. Variants were called using freebayes/ 1.0.2-6-g3ce827d (Garrison and Marth 2012) with modified arguments (-- pooled-discrete --pooled-continuous --report-genotype-likelihood-max --allele-balance-priors-off --min-alternate-fraction 0.1) and LoFreq/2.1.2 (Wilm *et al*. 2012) in a paired mode with their genetic ancestor. Called variants were filtered for uniqueness against their genetic ancestor(s) using bedtools/2.26.0. The variants were filtered for quality using bcftools/1.9 (Table S3). The filtered variants were annotated (Pashkova *et al*. 2013) and manually inspected for accuracy using the Interactive Genomics Viewer (IGV) (Robinson *et al*. 2011).

### Copy Number and Rearrangement Analysis

Using 1000 base pair sliding windows (IGVtools), normalized by the mean total read depth across the genome (GATK/3.7), the copy number was plotted and manually inspected for changes in copy number with the sample’s genetic ancestor. Copy number change breakpoints were manually inspected to determine the type of rearrangement using split and discordant reads, generated with BWA-mem (Li 2013), SAMBLASTER/0.1.24 (Faust and Hall 2014), and SAMtools.

### Data and Reagents Availability

All whole genome sequence data have been deposited at https://www.ncbi.nlm.nih.gov/sra with the bioproject accession number PRJNA544115 and will be made publicly available upon publication of the article. All other data necessary for confirming the conclusions of this article are fully within the article, its tables and figures or in supplementary material uploaded to GSA figshare. Yeast strains are available upon request.

## Results

### Altering the replication origins in the yeast rDNA locus using CRISPR/Cas9

Each repeat in the rDNA locus of *S. cerevisiae* contains a potential origin of replication between the divergently transcribed 35S and 5S genes in a region designated *NTS2* (non-transcribed spacer 2; Figure 1A). This origin sequence is inefficient in its native location (in many of the repeats the origin does not participate in replication initiation; Brewer and Fangman 1988; Muller *et al*. 2000; Pasero *et al*. 2002) and plasmids that contain the rDNA origin as their autonomous replication sequence (ARS) are poorly maintained (Larionov *et al*. 1984; Miller *et al*. 1999; Kwan *et al*. 2013). To determine whether the origin is essential for any of the diverse rDNA functions, we set out to use the CRISPR/Cas9 system to edit the entire set of repeats (see Methods and Materials; Figure S1A). We replaced the endogenous 11 bp A+T-rich ARS consensus sequence (ACS; Figure 1B; Figure S1B, Table S1) with a G+C block that contains four potential NGG PAM site alterations. We called these transformants rARS^GC^−1 and −2 and refer to their rDNA origins as *rARS^GC^* (Figures S2 and S3A). We also characterized two deletions that removed the origin region including all guide RNA sites (Figures S2 and S3B; Table S1). We called these strains rARS^Δ^−1 and −2 and refer to their rDNA origins as *rARS^Δ^*.

To introduce alternate origins into the rDNA, we repeated the editing experiments, this time either introducing non-rDNA origins or re-introducing the wild type rDNA origin. We chose two versions of the chr IV origin *ARS1* to replace the rDNA origin—a 100 bp fragment of the wild type version of *ARS1* and a multiply substituted version of *ARS1*, called *ARS1^max^*, that contains approximately 50 mutations discovered through a deep mutational scan, each of which contributed to improved plasmid maintenance (Liachko *et al*. 2013; Figure S1B; Table S1). We refer to these strains as rARS1-# and rARS1^max^-#. As a control for editing efficiency, we also reintroduced the wild type rDNA origin and the four PAM site mutations. We refer to these strains with the edited *rARS^“wt”^* as rARS^“wt”^−2, rARS^“wt”^−3 and rARS^“wt”^−12. Based on molecular analyses of the transformant clones (Figures S2, S3 and S4) we conclude that the editing of the entire set of rDNA origins is both efficient and complete.

### Assessment of origin function in strains with modified rDNA origins

To confirm that the modifications to the rARSs had altered origin function, we collected asynchronously growing cultures to analyze replication intermediates by 2D gel electrophoresis (Brewer and Fangman 1987). We digested chromosomal DNA with NheI to examine a ~4.5 kb fragment that contains the rDNA origin near its center (Figure 1A). We hybridized the Southern blots of the 2D gels with the *NTS2* probe looking for evidence of bubble structures that would signal initiation within the fragment (Figure 2). BY4741 produces the standard pattern of bubbles and Ys in a ratio of approximately 1:1.5, a ratio that indicates that approximately one out of every 2 to 3 repeats has an active origin and the other adjacent repeats are passively replicated by a fork from the upstream active origin. We find no bubbles among the replication intermediates for the rDNA loci in rARS^GC^−1 and rARS^Δ^−1, only Ys that are produced through passive replication (Figure 2).

**Figure 2:**
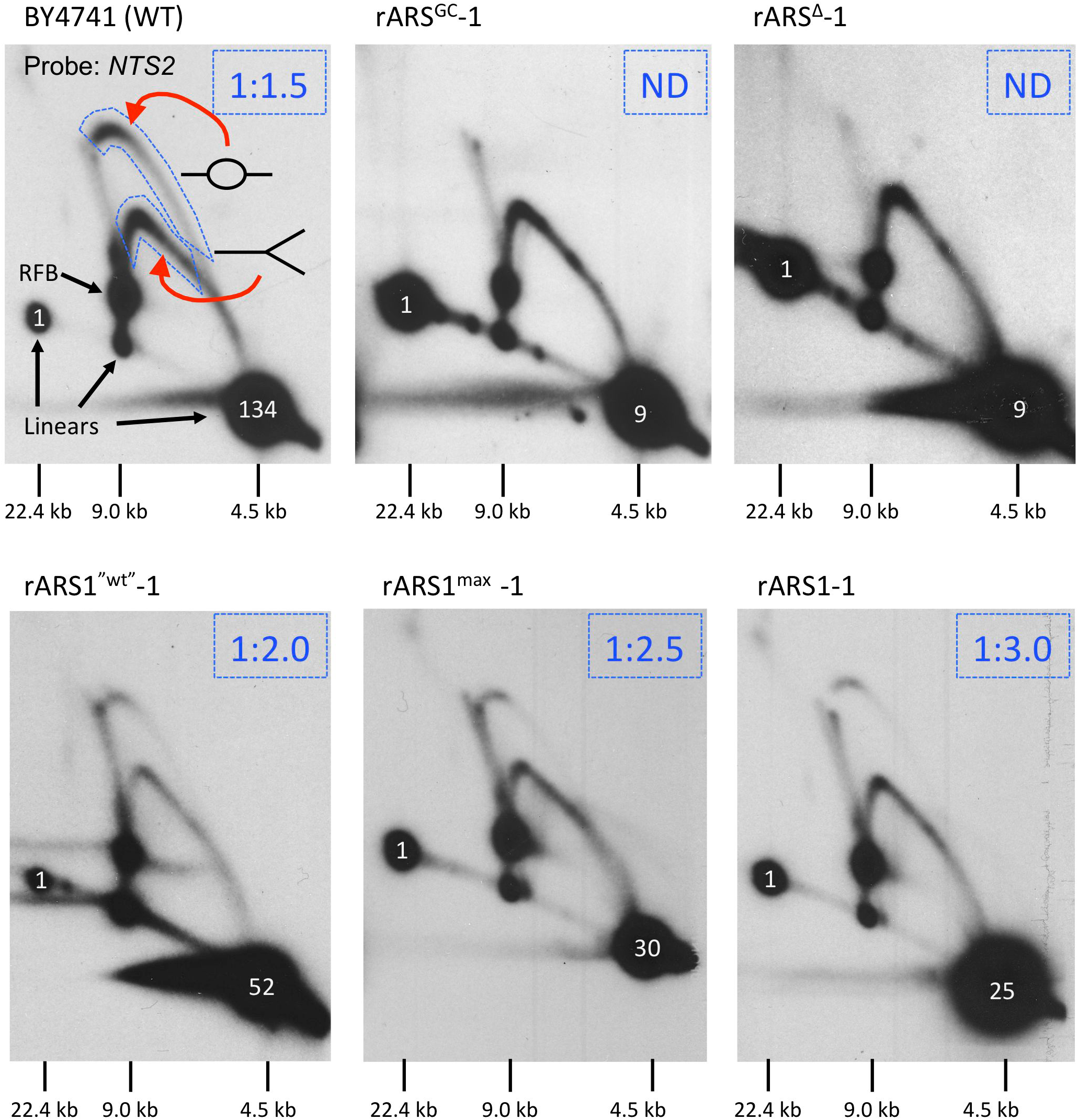
Assessing origin function in the rDNA locus by 2D gel electrophoresis. DNA isolated in agarose plugs from asynchronously growing wild type and edited strains was cleaved with NheI to release the *NTS2* fragment on a 4.5 kb fragment. Bubble-shaped molecules, indicative of initiation, are more retarded in the second dimension than Y-shaped molecules. We assessed Initiation efficiency by comparing the intensity of the bubble arc relative to the Y arc (areas surrounded by dashed lines). The bubble:Y ratios are indicated in the upper right corner of each 2D gels. ND indicates no signal above background was measured for the bubble region. Sizes of relevant linear species are indicated below each 2D gel panel. Numbers in white superimposed on the 4.5 kb fragments indicate the copy number estimated from the single copy sequence at 22.4 kb.

We conclude that the GC replacement and the deletion of the rARS reduce initiation efficiency in the rDNA below the level of detection by this assay. 2D gel analysis of strains with *rARS^“wt”^*, *ARS1^max^* and *ARS1* replacements show that these origins are functional after reintroduction into *NTS2* although the levels of replication initiation were somewhat lower than for the parent BY4741 strain—one out of every 3 to 4 repeats has an active origin (Figure 2).

A second feature revealed by the 2D gel analysis was a drastic change in copy number of the rDNA after origin editing. The telomere-proximal junction between rDNA and unique sequences produces a 22.4 kb NheI fragment that also contains the *NTS2* sequence (Figure 1A). By comparing the intensity of hybridization of the 4.5 kb *NTS2* linear fragment to this junction fragment, we determined that this particular isolate of BY4741 has ~134 rDNA repeats, while both rARS^GC^−1 and rARS^Δ^−1 have ~9 copies (Figure 2). Including a functional ARS on the repair template produced transformants with higher repeat numbers, but significantly lower than the BY4741 parent (Figure 2).

### Phenotypic properties of strains with modified rDNA origins

rARS^GC^−1 and rARS^Δ^−1 were slow to come up on the transformation plates and maintained their slow growth upon restreaking (Figure S2B). They were still able to utilize glycerol as the sole carbon source, ruling out a respiratory defect as the cause of slow growth. The same was true for all of the transformants that yielded smaller or no *NTS2* PCR fragments. Those transformants that escaped editing by Cas9 grew significantly faster (Figure S2B), and by avoiding the largest colonies on transformation plates in subsequent experiments we were able to bias recovery of edited clones. In liquid synthetic complete medium *rARS^GC^* and *rARS^Δ^* variants maintained their slow growth. In comparison to the 90-100 minute doubling time for BY4741, rARS^Δ^−1 had a doubling time of ~195 minutes and rARS^GC^−1 grew with a slightly longer double time of ~210 minutes (for examples, see Figure 4A). To explain the slow growth phenotypes of the modified clones we entertained several hypotheses, which are neither exhaustive nor mutually exclusive: 1) Cleavage by Cas9 resulted in a broken chr XII, which delayed cell division. The failure to detect wild type PCR product or bubbles on 2D gels would be explained by this persistent cleavage. 2) Cas9 cleavage resulted in a large-scale reduction of the rDNA copy number as excised rDNA repeats were unable to replicate or segregate efficiently and were eventually lost by the cells. The reduction in rDNA template would compromise ribosome production and result in slow growth. 3) Without functional origins of replication, the completion of chromosome XII replication was delayed as a fork from the adjoining, upstream unique origin would have to traverse the entire rDNA locus. The extension in S phase would delay the cell cycle.

To assess the possible causes of the slower growth we analyzed whole chromosomes from the transformants by <u>c</u>ontour-clamped <u>h</u>omogeneous <u>e</u>lectric field (CHEF) gel electrophoresis using conditions that allowed us to focus on chr XII and the rDNA locus. We looked for broken chr XII, extrachromosomal rDNA fragments, or delayed replication of chr XII as evidenced by a failure of chr XII to enter the gel. In the ethidium bromide stain of the CHEF gels (Figure 3A; compare with Figure S5A), all 16 of the slow growing clones appeared to be missing their largest chromosome (chr XII). However, hybridization of the Southern blot with the proximal (*CEN12*) and distal (*MAS1*) probes (Figure 1A) indicated that each of these slow growing strains had a shortened version of chr XII that co-migrated with chromosomes VII and XV (~1090 kb; Figure 3B; compare to Figure S5 B and C). Longer exposures revealed two different sub-chr XII fragments that are consistent with the size of the two chromosome halves if cleavage within the rDNA were occurring (~460 kb for the *CEN12* chromosome arm and ~600 kb for the distal arm containing *MAS1*; Figures 3C and D). (The two fragments represent ~5% of the chr XII signal.) To determine whether additional rDNA repeats had integrated elsewhere in the genome or were persisting as extrachromosomal molecules, we reprobed the blots with an *NTS2* rDNA probe (Figure S6). We did not detect free rDNA repeats nor repeats that were housed on other chromosomes. We also did not detect disproportionate hybridization of rDNA to the wells of the slow-growing clones, indicating that chr XII from these stationary phase samples had completed replication.

**Figure 3:**
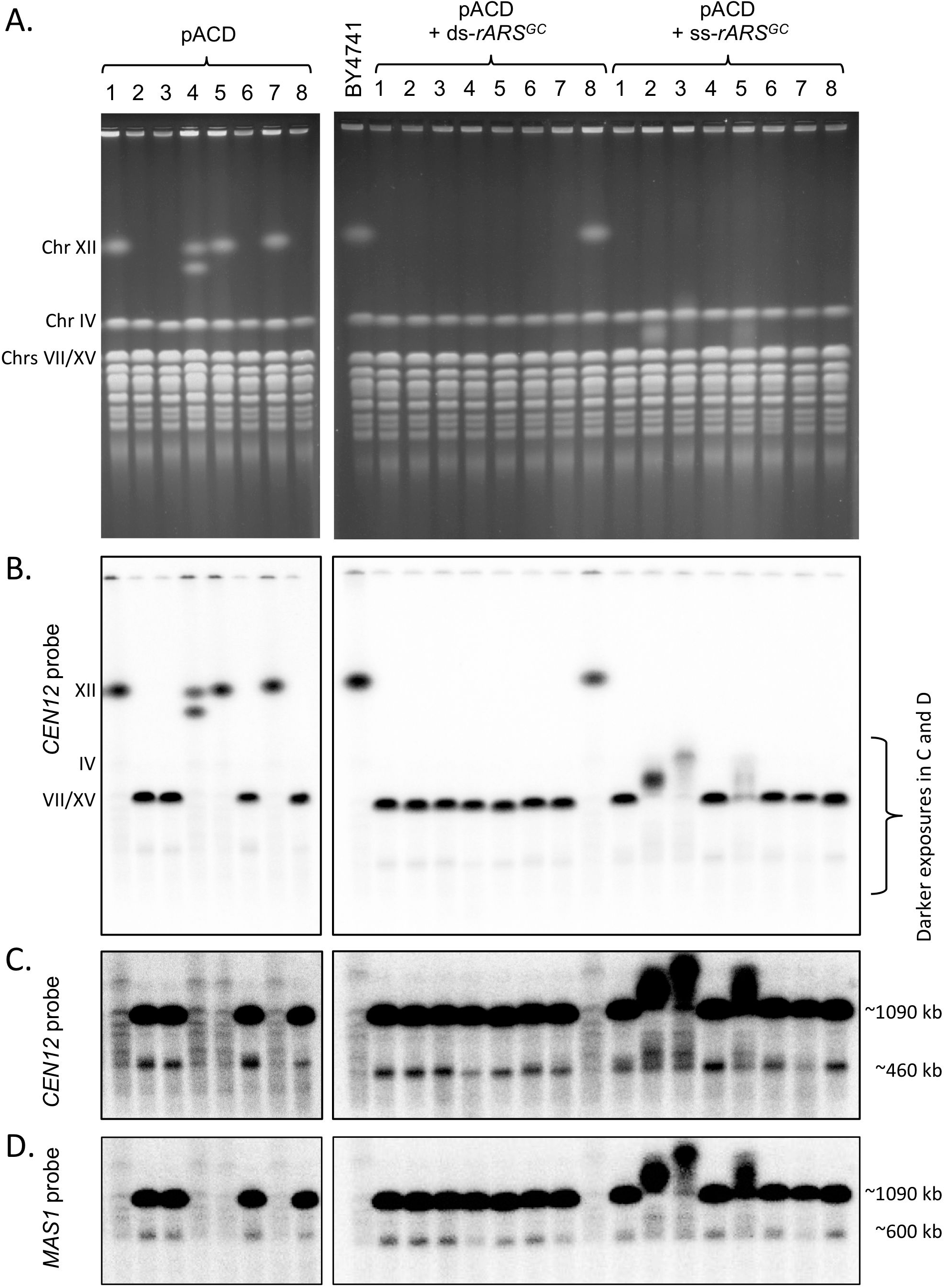
CHEF gels analysis of CRISPR/Cas9 transformants. A) The ethidium bromide stained CHEF gels, run under conditions that maximize the separation of Chr XII, are shown for the same three sets of eight strains from Supplemental Figure 2. B) Southern blots of the CHEF gels were probed with a fragment from the centromere region of chr XII. The small chr XII present in the majority of samples is co-migrating with chrs VII and XV. C) A longer exposer of the bracketed section of the blots shown in (B) reveals a sub-chromosomal fragment of ~450 kb. D) The blots were stripped and reprobed with *MAS1*, which lies on the distal side of the rDNA. A subfragment of ~600 kb is found among the same transformants that had the ~450 kb fragment.

Because it seemed unlikely that the 5% of cells with a broken chr XII could account for the roughly 2.3-fold increase in doubling time we turned to the hypothesis that the reduction in rDNA copy number (hypothesis #2 above) was the chief cause of slow growth. Since unique sequences make up ~1060 kb of chr XII the number of rDNA repeats following CRISPR/Cas9 editing could be as low as 3 or 4, although the resolution using these running conditions is not adequate to get an accurate estimate of rDNA length. The CHEF gel of the rDNA^“wt”^ strains demonstrates that they also have a reduced rDNA copy number (Figure S7A). To estimate the rDNA copy number of the rARS^“wt”^ and rARS^Δ^ clones, we digested genomic DNA in gel plugs with FspI to release the entire rDNA locus as a single restriction fragment (Figure 1A). Using the wild type chromosomes as size references, we estimate that the three *rARS^“wt”^* clones have between 25 and 75 repeats, each clone having broad distributions of ±10 repeats (Figure S7B). The set of uncharacterized deletion strains from this transformation have in the range of 6 to 11 rDNA copies, a similar value to that of rARS^GC^−1 and rARS^Δ^−1.

We suspected that the slow growth of rARS^GC^−1 and rARS^Δ^−1 was the result of reduced rRNA production since these strains had just nine copies of the rDNA repeats. To measure the relative rRNA content in rARS^GC^−1, we quantified the amount of 25S ribosomal RNA relative to the DNA of a single copy nuclear gene (*ACT1*) and found a 35% reduction in 25S rRNA in rARS^GC^−1 relative to the parent strain BY4741 (Figure S8). We conclude that hypothesis #2 most easily explains the slowed growth of the edited rDNA strains: Cas9 cleavage releases most of the rDNA repeats from chr XII and these repeats are subsequently lost from the cell. This situation leads to a drastically reduced chr XII size and rDNA repeat copy number, which compromises rRNA production, resulting in slowed growth. Crossing rARS^GC^−1 or rARS^Δ^−1 to BY4742, a wild type strain with an unedited rDNA array, alleviated the slow growth phenotype, suggesting that the wild type chromosome with its ~150 rDNA repeats was meeting the demand for ribosomes. However, in the heterozygous diploids the mutant rDNA locus further contracted to ~2-5 copies (Figure S7C), suggesting that replication delay of the large origin-free, ~10 copy mutant rDNA locus also may have contributed to the slow growth of the haploids.

### Re-expansion of the rDNA locus after origin replacement

We estimate that transformants had grown for 40-45 generations by the time we could examine the size of chr XII on CHEF gels or examine replication intermediates on 2D gels. Yet, at a similar number of generations the transformants with different repair templates had achieved different rDNA repeat sizes (Figure 2 and Figure S7B). To determine whether the differences in chr XII sizes and initiation efficiency among the *rARS^“wt”^*, *rARS1^max^* and *rARS^GC^* or *rARS^Δ^* strains were due to differences in amplification rates for the edited repeats, we carried out long-term growth experiments using one or more of three different culturing methods: daily serial passaging to keep the cultures in log phase growth, daily dilution of 1:1000 letting the cells reach stationary phase each day, or continuous culture in a turbidostat where the optical density was held constant. For both the *rARS^“wt”^* and *rARS1^max^* strains (passaged daily after reaching stationary phase) we found an initial rapid increase in rDNA copy number followed by a steady, modest increase in chr XII of approximately 2 repeats per 10 generations of growth that failed to reach the size of the original copy number of ~150 rDNA copies after an additional ~100 generations (Figure S9A and B).

We examined strains rARS^GC^−1 and rARS^Δ^−1 that we kept in log phase growth over a period of 13 days, monitoring optical density daily to assess growth rates. The cumulative ODs of two individual colonies of rARS^GC^−1 and three colonies of rARS^Δ^−1 (Figure 4A and B) show a striking difference: approximately 50 generations (160 hrs; dotted line Figure 4A) into the experiment, both colonies of rARS^GC^−1 began growing more rapidly and continued growing at the increased rate for the remainder of the experiment. rARS^Δ^−1 colonies did not show this striking change in growth properties. The CHEF gel of rARS^Δ^−1 clones shows that chr XII is unable to expand its rDNA repeats significantly, while the rARS^GC^−1 clones begin showing signs of expansion at the same time that the cultures began growing faster (Figure 4C). At ~50 generations the cells were heterogeneous with a majority of the chromosomes remaining at the initial size of ~10 repeats but by the end of the experiment, all cells had expanded their rDNA loci to a median copy number of ~25 repeats. We saw no increase in the size of chromosome XII in the three clones of rARS^Δ^−1 (Figure 4D).

**Figure 4:**
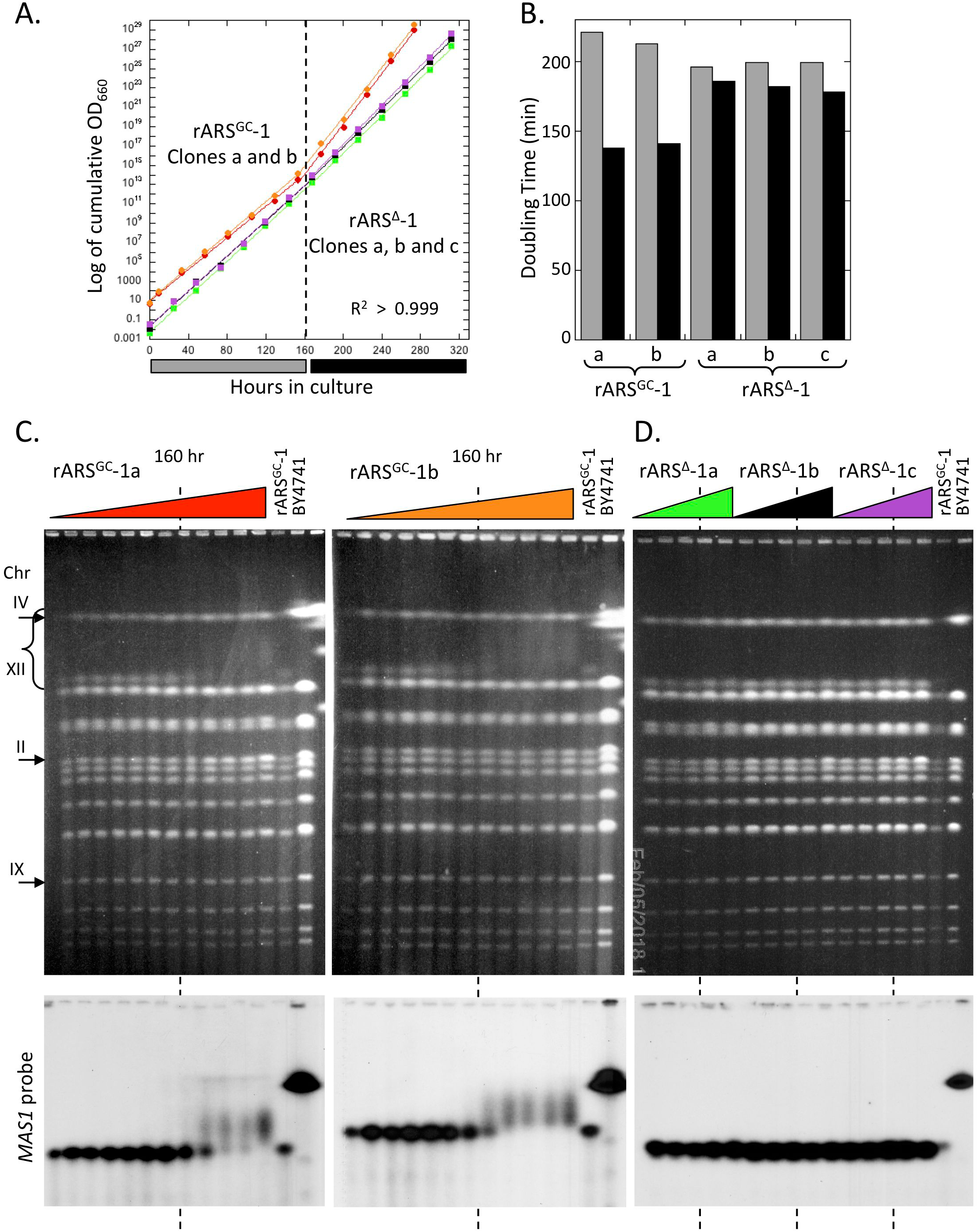
Monitoring changes to growth rates and chr XII size during serial passage of rARS^GC^−1 and rARS^Δ^−1. A) Cumulative OD for cultures kept in continuous logarithmic growth. Two clones of rARS^GC^−1 (−1a and −1b) and three clones of rARS^Δ^−1 (−1a, −1b and −1c) were followed for ~13 days. B) Population doubling times for the first (grey) and second (black) halves of the continuous growth experiments in A. C) and D) CHEF gel analysis of the changes to chr XII size over ~13 days: gels in C contain samples collected each day, gels in D contain samples collected on days 0, 3, 6, 9 and 12. The conditions for CHEF gel electrophoresis allow the detection of subtle changes in chr XII size. Only the top halves of the Southern blots probed with *MAS1* are shown.

We repeated long-term growth of *rARS^GC^* and *rARS^Δ^* strains using turbidostats. These continuous culture devices detect when faster growing variants sweep the population because they trigger an increase in the delivery of medium to cultures in an attempt to maintain a constant culture OD. We analyzed five clones of *rARS^GC^* and three clones of *rARS^Δ^* in turbidostats for a period of ~7-8 days (or 85-100 generations) and for all but one of them we find evidence for increased growth rates by the end of the experiment—some more modest than others (Figure 5A; Figure S10A). We collected samples once each day and prepared genomic DNA for CHEF gels (Figure 5B and C; Figure S10B and C). By the end of the experiments, we found increases in rDNA copy number for the *rARS^GC^* cultures but severely restricted amplification in the *rARS^Δ^* cultures—findings consistent with the logarithmic, serial transfer experiments.

**Figure 5:**
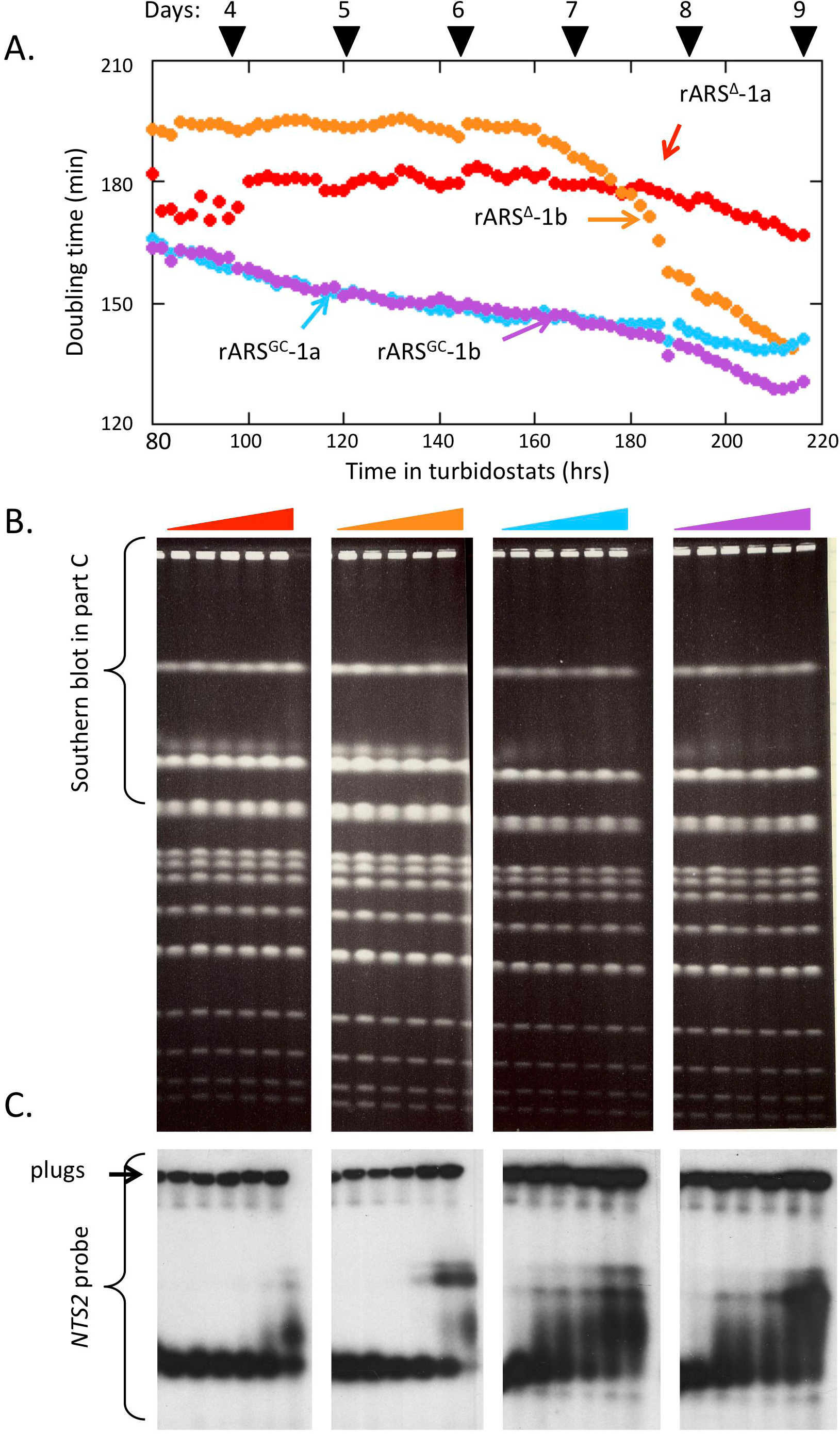
Selection for suppressors of slow growth in turbidostat cultures of *rARS^GC^* and *rARS^Δ^*. A) The growth rates calculated from the pump speeds for two colonies of *rARS^GC^* and *rARS^Δ^* over the last ~six days of the turbidostat runs. B) CHEF gels for samples harvested on the days indicated in (A) by the black triangles. C) Southern blot analysis of changes in chr XII/rDNA over the course of the continuous growth.

### Return of replication initiation to the rDNA *NTS2* region of *rARS^GC^* clones during rDNA expansion

We isolated single clones from both day 10 batch cultures of rARS^GC^−1a and −1b (Figure 4) to reanalyze rDNA origin firing after ~100 generations in continuous log phase. 2D gel analyses of both clones show low-level initiation within the *NTS2* region and quantification of the spots corresponding to the 22.4 and 4.5 kb linear fragments confirms the expansion of rDNA copy number (Figure 6). To explain the reappearance of bubbles in the rDNA we entertained three possibilities that could explain the return of origin activity to the rDNA *NTS2* region that was allowing the re-expansion in repeat number beyond the ~10-13 copy ceiling: 1) The initial clone had retained a single wild type copy of the *rARS* and that, over continuous selection for rapid growth, cells had used this single copy to convert the *rARS^GC^* mutation back to the wild type sequence. 2) Other sequences in the *NTS2* had mutated to create a new *rARS*. 3) A mutation arose elsewhere in the genome in a gene whose altered product was able to recognize *rARS^GC^* or some other sequence in NTS2. However, as described below, we found no evidence for any of these hypotheses.

**Figure 6:**
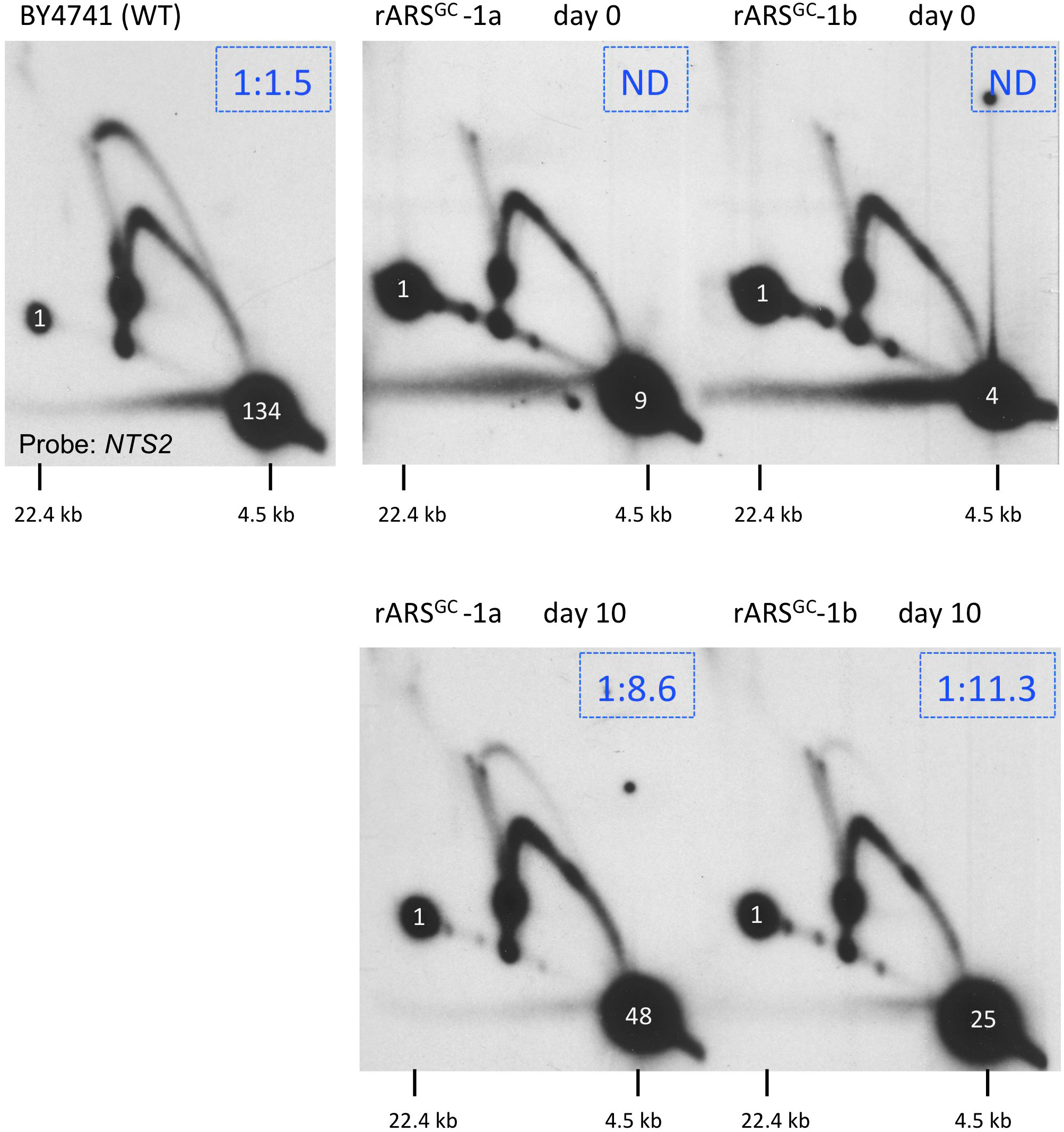
2D gel analysis of rARS^GC^−1a and rARS^GC^−1b at day 0 and single clones from these cultures from day 10 of the continuous growth experiments. The Southern blots were probed with the *NTS2* probe to assess origin function. rDNA copy number estimates (in white type) are deduced from the relative signal of the 4.5 kb fragment to the 22.4 kb junction fragment. The BY4741 and rARS^GC^−1a day 0 gels are the same as in Figure 2.

To look for return of the wild type rARS sequence we PCR-amplified and Sanger-sequenced a ~600 bp fragment containing the edited regions. We found that the sequences from the faster-growing clones were identical to the initial edited sequence: the *rARS^GC^* replacement and PAM mutations were still present and there was no trace of the original wild type sequence (Figure S3).

To look for origin activity in the flanking regions, we PCR amplified the entire *NTS2* region, cloned it into a non-replicating plasmid, and checked the library of plasmids for any that were able to replicate autonomously in yeast. The few transformants we recovered had all integrated the selectable marker into the genome, indicating that there were no *NTS2* regions that had acquired a new origin capable of maintaining a plasmid.

To test the hypothesis that a trans-acting mutation elsewhere in the genome was recognizing the altered *rARS*, we used a clone from day 10 of the evolution as a host in a transformation experiment in which we introduced the *rARS^GC^* plasmid library (see above). Again, we recovered no transformants that were able to support replication of the *rARS^GC^* plasmid library members as free plasmids, arguing against the possibility that a second-site mutation was allowing initiation within the edited rDNA.

From these combined results we have no direct result supporting a genetic cause for the return of initiation to the rDNA *NTS2* region. Yet, the striking difference in rDNA expansion between the rARS^GC^−1a and −1b clones and the rARS***^Δ^***−1a −1b, and −1c clones suggested that the sequence of the *NTS2* region was somehow important for the amplification of the rDNA and for the generation of faster growing suppressors. To test this conclusion further, we sequenced the genomic DNA from day 10 populations and clones of the rARS^GC^−1a and −1b growth experiments. In comparison to the day 0 clones of these two cultures, there were no alterations to the rDNA repeats and no SNPs in genes relevant to DNA replication (Table S2). However, it was notable that rARS^GC^−1 had indels in *IRA1* or *IRA2*, which may have been important for enhancing the growth and subsequent improved fitness of the evolved strains derived from this background (Colombo *et al*. 2004)

### Chromosome aneuploidy contributes to improved growth of *rARS^GC^* and *rARS^Δ^* strains

The lack of any single nucleotide variants that could explain improved growth rates led us to investigate large-scale changes by using the WGS data to investigate copy number differences across the genome. Read-depth analysis of the WGS of the day 10 clone of rARS^GC^−1a from the serial passaging experiment (Figure 4) suggested that the cells had become disomic for chr II (Figure 7A). Consistent with this idea, the ethidium bromide photograph of the CHEF gel for rARS^GC^−1 (Figure 4C) suggested that by the end of the experiment the intensity of chr II signal was elevated relative to that of the surrounding chromosomes. Comparative hybridization with probes from the centromere-adjacent regions of chr II and chr IX confirmed the chr II disomy for rARS^GC^−1a (Figure S11). The CHEF gel of the three *rARS^Δ^* evolutions (Figure 4D) indicated that in two of them (rARS^Δ^−1b and −1c) there had also been an amplification of chr II.

**Figure 7:**
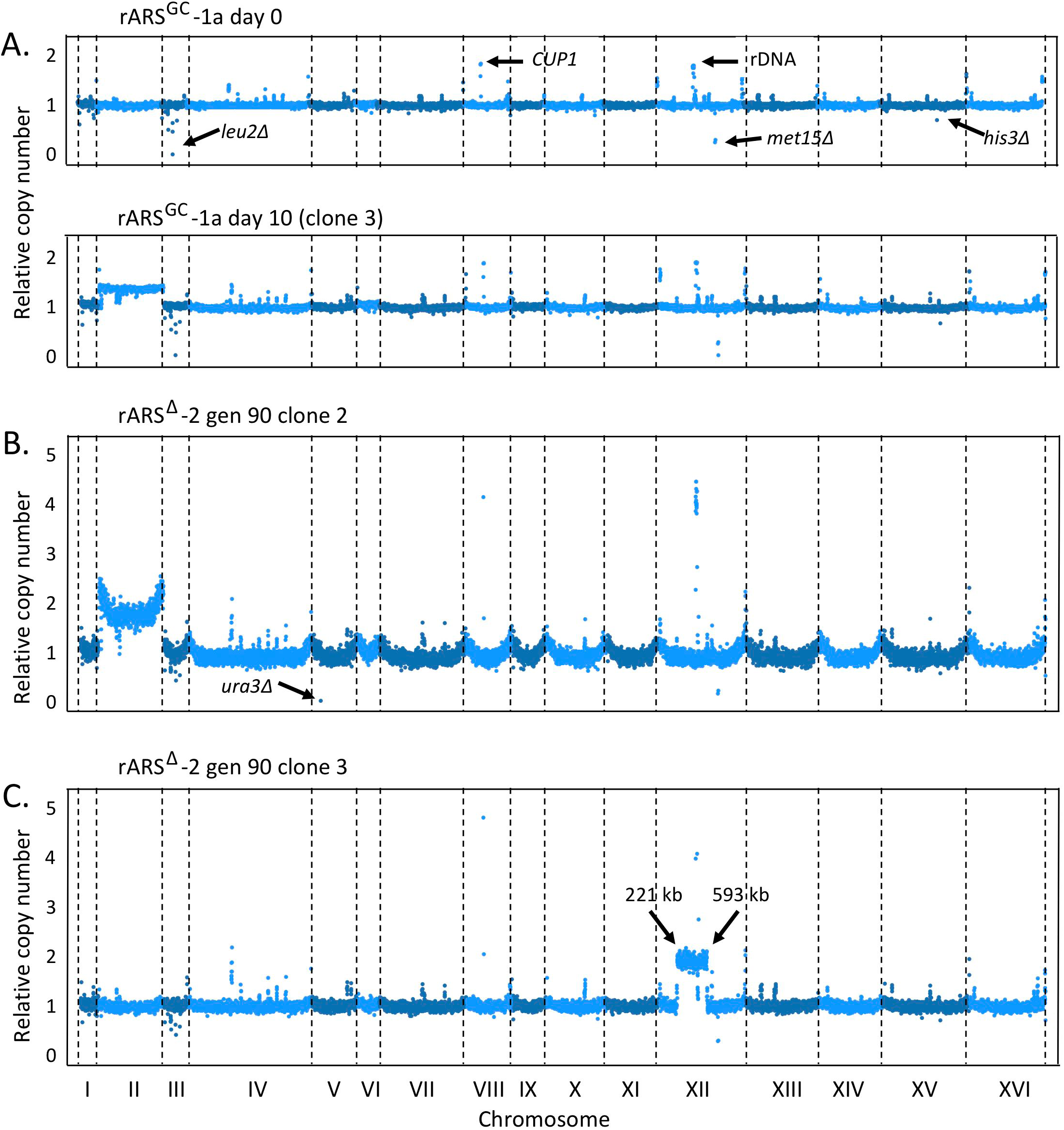
Read-depth analysis from whole genome sequencing (WGS) for three evolved *rARS^GC^* clones. A) Genomic DNAs from rARS^GC^−1a on day 0 and a clone from day 10 of the continuous passaging experiment were sequenced by WGS. Read-depth measurements across 1 kb bins were normalized to the genome average. Individual values below ~1.0 are seen for the various auxotrophic deletions in the parent strain BY4741 (*leu2Δ*, *ura3Δ*, *met15Δ* and *his3Δ*). Individual values above ~1.0 indicate multiple mapping of Ty transposable elements, the repeated *CUP1* locus on chr VIII, and the rDNA locus on chr XII. In this clone from day 10, half of the cells appear to be disomic for chr II. B) and C) WGS read-depth analysis for two clones of rARS^Δ^−2 isolated on day 9 (90 generations) of the continuously passaged culture. Clone 2 is disomic for chr II. (The increased copy number of sequences near telomeres is an artifact of the library preparation.) Clone 3 has a segmental aneusomy for a region of chr XII between two directly repeated Ty long terminal repeats at positions 221 and 593 kb.

As it appeared that chr II aneuploidy was a recurring event that we detected following the reduction in rDNA replication initiation or copy number, we examined the serial passaging experiments with the independent isolates of *rARS^GC^* and *rARS^Δ^* (clones rARS^GC^−2 and rARS^Δ^−2). We found comparable results for rDNA amplification: there was no significant amplification of the rDNA locus in rARS^Δ^−2 but slight amplification in rARS^GC^−2 (Figure S12A and B). From these two experiments we isolated four individual clones at generation 90 for further analysis (Figure S12A). We compared Cen2/Cen9 hybridization to track and quantify the appearance of the chr II disome over time (Figure S12 C, D and E). The population of rARS^GC^−2 was essentially homogeneous for chr II disomy by generation 50 and all four clones from day 9 of the passaging experiments were disomic for chr II. No significant increase in chr II was observed during passaging of rARS^Δ^−2; however, clone#2 from generation-90 contained an additional copy of chr II (Figure S12E).

To determine whether disomy for chr II actually provides a selective advantage for the strains with reduced rDNA copies, we crossed clone 1 from the rARS^GC^−2 passaging experiments (Figure S12A and E) with BY4742 and examined the growth properties of segregants containing the parental and derived chr XIIs with and without an extra copy of chr II (Figure 8A). We examined six complete tetrads for their patterns of chromosome segregation and analyzed the growth properties for the spores of four of these tetrads (Figure 8B and C). The presence of an extra copy of chr II improved the growth of the strains that contained *rARS^GC^* by nearly 30 minutes (~16% improvement in doubling time) but increased the doubling time by 8 minutes (~10%) in strains with the wild type rDNA origin and intact rDNA array.

**Figure 8:**
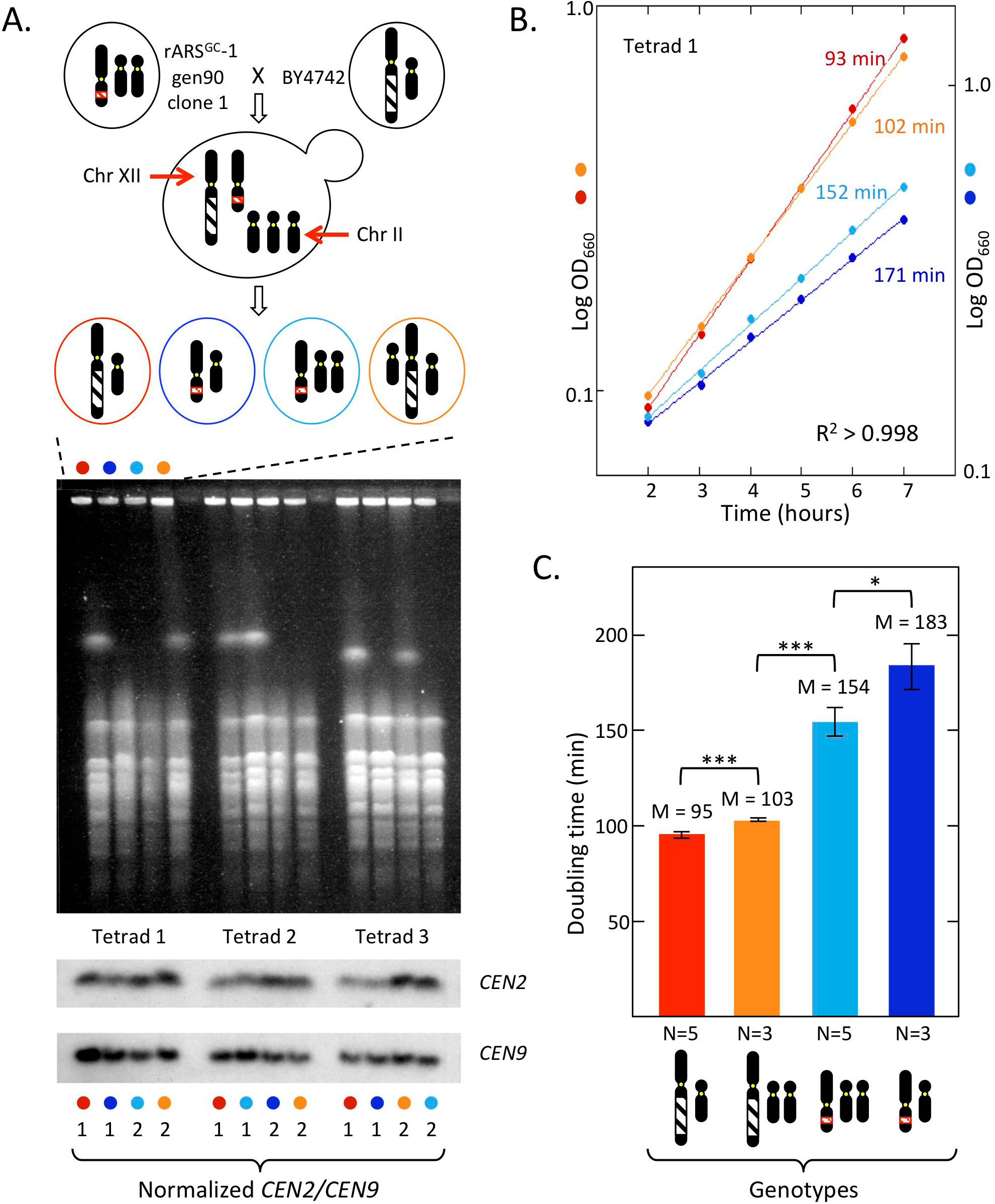
Scheme for analyzing the growth properties of chr II disomy in strains with the native rDNA and with *rARS^GC^*. A) Top: the scheme for mating clone 1 from gen-90 of the rARS^GC^−2 serial passage experiment (Supplemental Figure 12A) with BY4742 and segregation to form a tetratype tetrad. Middle: Ethidium bromide stained CHEF gel of three tetratype tetrads from the cross. Bottom: Southern blots to follow segregation of chr II disome. B) Growth rates for the four spores from Tetrad 1 in part A. C) Average growth rates for similar genotypes isolated from four independent tetrads. Significance values were calculated using student T-test: P < 0.05 (*); P < 0.005 (***).

We performed WGS for the four generation-90 clones from rARS^Δ^−2 (Figure S12A). Read depth analysis confirmed the chr II disomy of clone 2 and revealed a different aneuploidy for clone 3 (Figure 7B and C). In this clone, a region of chr XII of ~ 370 kb, from coordinate 221 kb (YLRWdelta 4) to 693 kb (YLRWdelta10) was present in two copies. Between these two long terminal repeats lie the rDNA locus and both *MAS1* and *CDC45*. This duplicated segment is located on chr XII and, along with ~10 copies of the rDNA repeats, accounts for the increased size of chr XII seen on the CHEF gel (clone #3; Figure S12C). The ratio of *MAS1* or *CDC45* to *CEN9* supports our conclusion that an intrachromosomal duplication between two LTRs has allowed the total copy number of rDNA repeats to double in this particular clone, without expanding either rDNA locus beyond the apparent limit of ~10 ARS-less repeats. Although we were unable to find a clone with this chr XII duplication from either of the rARS^GC^−1 passaging experiments, we did see the transient appearance of this larger chr XII during the later days of the rARS^GC^−1a passaging experiment (Figure 4C) and in many of the turbidostat cultures (Figure 5C and Figure S10C). We also sequenced the population samples from the last day of each of the eight turbidostat cultures (Figure 5; Figures S10 and S13; Tables S2 and S4). For all eight of the cultures we find chromosome aneuploidies/aneusomies that accumulate to different extents in the populations—five cases of chr IV disomy, two cases of chr XII disomy and one case of an internal duplication of the rDNA locus on Chr XII (Figure S13; Table S4). For none of the populations do we find common single nucleotide variants that could easily explain the entirety of the improved growth rates (Table S2).

### The effect of rDNA origin replacement on extrachromosomal rDNA circles (ERCs) and rDNA re-expansion

Using the hygromycin system of rDNA manipulation, Ganley et al. (2009) introduced two different versions of the *ARS1* origin, along with a selectable *URA3* marker, into the yeast rDNA locus that had been reduced to two copies. They found that the efficiency of the rDNA origin influences both the production of ERCs and the rate that the rDNA locus re-expands. ERCs are thought to be formed primarily through homologous recombination between adjacent or nearby rDNA repeat units, which results in excision of a circular DNA molecule containing one or more complete rDNA units. ERCs are capable of self-replication based on the strength of the *rARS* and are asymmetrically retained in the mother cell, causing them to increase dramatically in copy number with mother cell age (Sinclair and Guarente 1997). To compare the two methods of altering the rDNA origin we analyzed our *rARS* modified strains for the presence of ERCs.

We anticipated finding differences in the steady state levels of ERCs among the edited strains, as the efficiency of the rDNA origin should affect their ability to propagate ERCs. For example, the *rARS^Δ^* strains might be able to produce ERCs by intramolecular recombination, but once formed they would not be able to initiate replication and would be diluted at cell division. To assess the relative ERC populations among different edited strains, we analyzed portions of freshly prepared plugs using conventional gel electrophoresis. Under these conditions, all of the yeast chromosomes run at limiting mobility with different circular multimers/topological forms running both above and below this band (Figure 9). By comparing the hybridization signals of the two lowest circular forms to that of the chromosomal rDNA band, we found that the steady state level of ERCs in these predominantly young cell populations is proportional to the potential efficiency of the different rDNA origins, with the *rARS1^max^* transformants maintaining >100 times higher levels of ERCs than BY4741. At the other extreme, we were unable to detect ERCs in either *rARS1^GC^* or *rARS^Δ^* strains.

**Figure 9:**
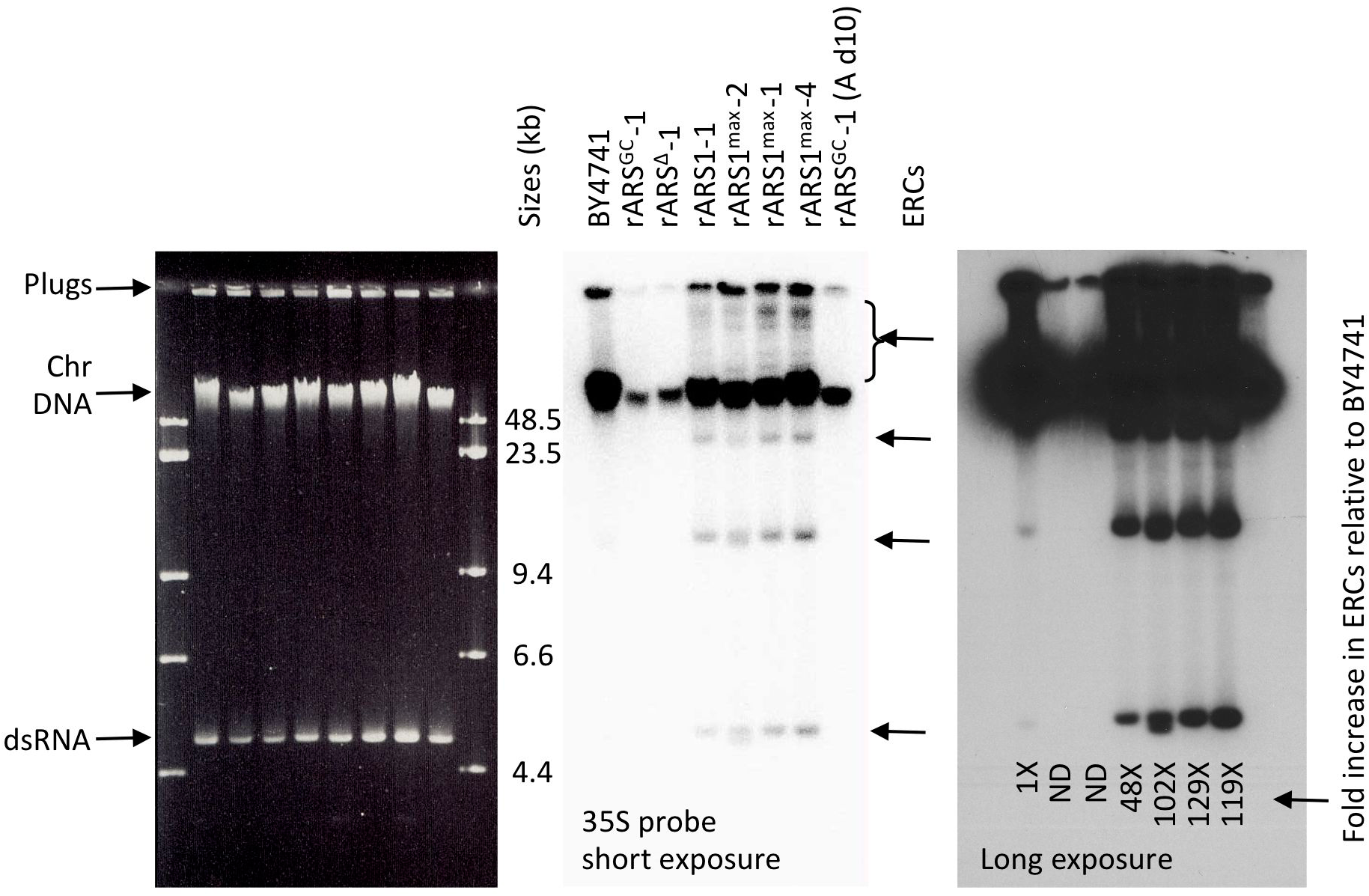
Comparison of ERC frequency among the different *rARS* replacement strains. The DNA in agarose plugs was run on a standard 0.5% agarose gel where chromosomes and circular molecules are easily distinguished. The Southern blot using a probe from the 37S gene is shown at two exposure levels. The ratio of hybridization to the two lower bands was quantified and compared to the signal from the chromosomal band. The abundance of ERCs in BY4741 was defined as 1 and other strains were normalized to that value. Because the edited strains all had lower chromosomal rDNA copy numbers, their estimates for ERCs on a per cell basis are underestimates.

The high level of ERCs in the *rARS1* and *rARS1^max^* strains prompted us to look more carefully at the initial transformants of both *rARS1* and *rARS1^max^* (samples at ~45 generations; Figure S14 A and B). Of the clones that had successfully replaced the *rARS*, there were five clones that had an even smaller chr XII than the rARS^GC^−1 parent clone (*rARS1^max^* clones 2, 4, and 7; *rARS1* clones 2 and 5; *MAS1* probe in Figure S14 A and B). On reprobing the blots with the 35S probe, we found that these clones showed little to no hybridization to chr XII, but additional rDNA-positive bands appeared at positions where no corresponding chromosome were present (horizontal arrows; Figure S14 A and B). These bands are consistent with the properties of circular molecules of different sizes (monomers, dimers, etc.) and topologies (supercoils and nicked circles) (Brewer *et al*. 2015). The suggestion that no *NTS2* sequences exist on chr XII is corroborated by the 2D gel analysis of NheI digested genomic DNA we performed on two of the clones: the *NTS2* probe no longer detected the 22.4 kb junction fragment between rDNA and the distal arm of chr XII (Figure S14 C). This extrachromosomal state of the rDNA is clonally stable on solid culture medium: we restreaked individual clones from the rARS1^max^−4 transformant and found, ~30 generations later, that each clone retained high levels of ERCs (Figure S15A). However, upon serial passaging of rARS1^max^−4 in liquid medium, the ERCs largely disappeared by 30 generations and chr XII again contained rDNA repeats (Figure S15B). Over the remaining ~100 generations the rDNA slowly expanded, but at a much slower rate than we observed for rARS1^max^−8 (compare Figure S15B with Figure S9B; Table S4).

### The consequence of rDNA origin replacement for replicative lifespan

A variety of observations in yeast and other eukaryotes have shown that the rDNA plays a significant role in determining replicative lifespan, which is defined as the number of daughter cells produced by a mother cell prior to irreversible cell cycle arrest (Mortimer and Johnston 1959). First, an *rARS* ACS variant found in the vineyard strain RM-11 reduced the efficiency of the rDNA origin and extended life span (Kwan *et al*. 2013). Second, a reduction in total ribosome levels through calorie restriction, chemical interventions such as rapamycin, or genetic manipulations through the deletion of ribosomal subunit proteins (Lin *et al*. 2000; Kaeberlein *et al*. 2005; Chiocchetti *et al*. 2007; He *et al*. 2018) all robustly extend lifespan across the evolutionary tree. Third, in yeast, the accumulation of ERCs in mother cells has been linked to lifespan (Sinclair and Guarente 1997). Deletion of Fob1, a protein that binds to the replication fork barrier (RFB) site in each rDNA repeat and blocks head-on collisions between RNA Pol1 and the replication complex, reduces ERC levels and dramatically extends lifespan (Defossez *et al*. 1999). As our edited rDNA strains have variations in their rDNA origins, significantly reduced rDNA copy number, reduced ribosomal RNA content (Figure S8), and variable levels of ERCs we decided to conduct replicative lifespan analysis on representatives of the different origin replacement strains we generated.

We micromanipulated individual, newly “born” daughter cells on agar plates. We monitored the cells, counting and removing consecutive daughter cells. BY4741 and the BY4741 strain with the rDNA locus from RM-11 (Kwan *et al*. 2013) were used as the normal (mean lifespan = 23 gen) and long-lived (mean lifespan = 41) controls, respectively. We anticipated that *rARS^GC^* and *rARS^Δ^* strains, with severely restricted origin activity in the rDNA, would have extended lifespans. Contrary to our expectations we found reduced lifespans for both strains (Figure 10 A and C), with *rARS^Δ^* having a median lifespan of only nine generations. Lifespans for the chromosomally integrated *rARS1* and *rARS1^max^* strains were similar to that of the *rARS^GC^* variants (Figure 10 B and C). Perhaps most surprising are the similar lifespans of the *rARS1^max^* variant that has only ERCs (rARS1^max^−4) and the *rARS^Δ^* strain which has no ERCs (Figure 10). Unfortunately we cannot currently measure either the rDNA copy number or the level of ERCs for the individual mother cells that were analyzed for lifespan. However, if these mother cells accurately represent the cultures from which they were isolated, then our data suggest that drastically reduced rDNA copy number masks any benefit to lifespan that may result from rDNA origin status or ERC levels.

**Figure 10:**
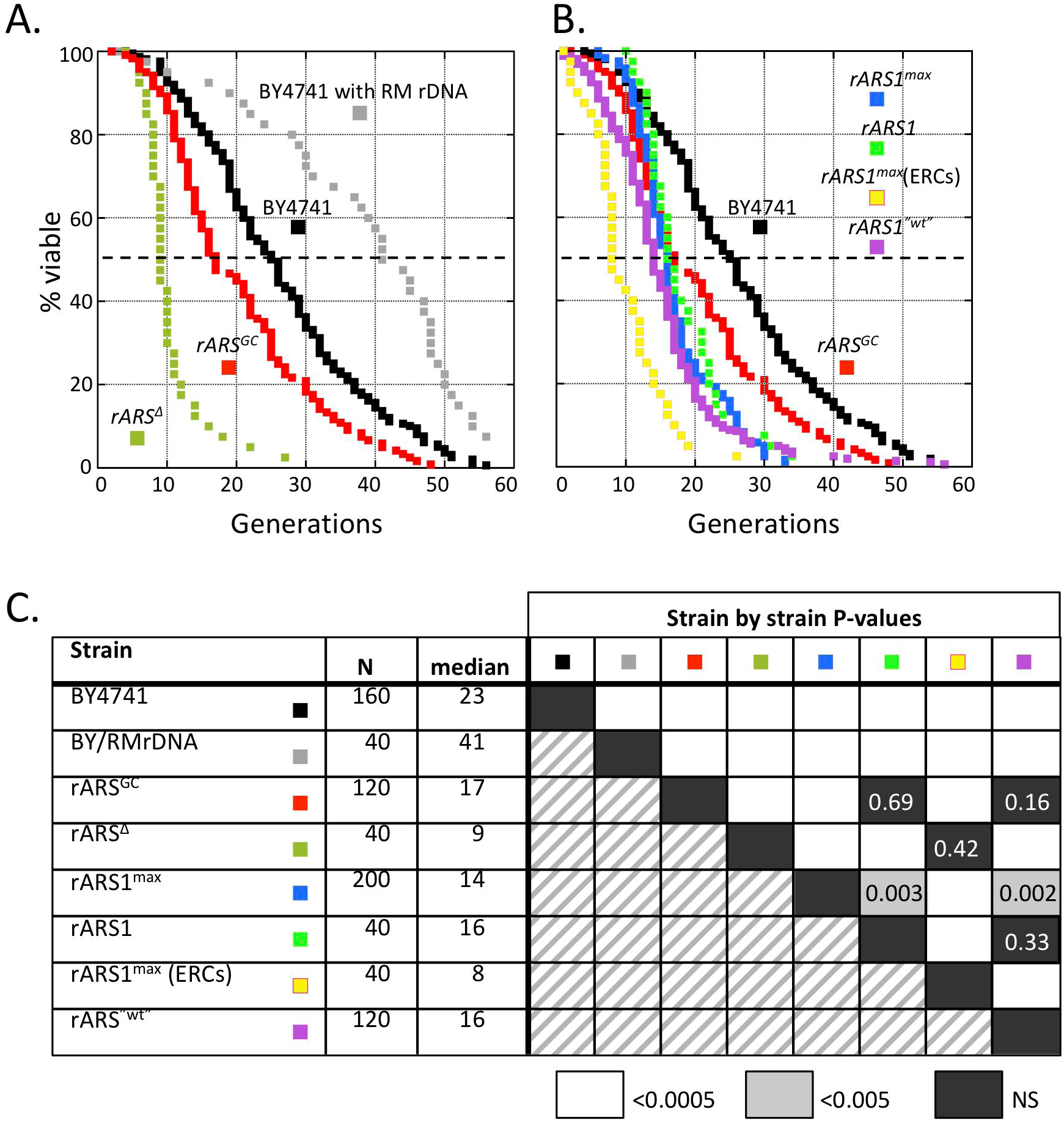
Replicative lifespans of rARS edited strains. A) Lifespan curves for the strains that were transformed with either *rARS^GC^* or *rARS^Δ^* compared with BY4741 and the long-lived BY strain with the rDNA from RM-11. B) Lifespan curves for strains transformed with functional ARSs. Multiple independent transformants were separately examined and the data pooled to generate the lifespan curves for *rARS1^max^* and *rARS^“wt”^*. The lifespan curves for *rARS1* and *rARS1^max^* (ERCs) were generated from single transformants. The BY4741 and *rARS^GC^* data are reproduced from A. C) The number of mother cells followed (N) and the median lifespans were determined for each class of mutant. P-values from the Wilcoxon Rank-Sum analyses for each pairwise strain comparison are shown. Highly significant values are indicated by white squares; non-significant (NS) values are indicated by black squares.

## Discussion

CRISPR/Cas9 and its many offshoots have been successfully used for editing of an enormous variety of genetic targets in many organisms. Targets have been almost exclusively single copy and worry about off-target cutting has been a prominent concern. In this work, we have set out to ask—can we edit tandem repeats, how efficiently and at what phenotypic costs? We have worked out conditions for targeting the ~150 identical targets in the rDNA locus simultaneously, to repair the breaks with templates of our choosing, and to follow the phenotypic consequences of the modified homogeneous rDNA arrays. From our WGS analysis, we find no obvious off-target edits (Table S2).

The rDNA locus in most organisms is difficult to genetically modify through more conventional techniques such as random mutagenesis or even targeted gene replacement strategies that require selectable markers because of the seemingly impossible task of altering every member of the array. Yet an ever-growing list of biological processes, in addition to ribosome biogenesis, are linked to the rDNA and the nucleolus that forms around the rDNA (Boisvert *et al*. 2007; Bahadori *et al*. 2018). Recently, rDNA copy number in several cancers have been found to be lower than in matched non-cancerous biopsies (Wang and Lemos 2017; Xu *et al*. 2017) and appear to be increased in some patients with schizophrenia (Chestkov *et al*. 2018). At this point, however, the causality of rDNA changes in these diseases is undetermined. In addition, variation in nucleolar size has been shown to correlate with lifespan (Tiku *et al*. 2017). In experimental organisms, sequence variations in the non-coding regions of the rDNA have been documented but again their significance is difficult to determine (Ganley and Kobayashi 2007; James *et al*. 2009; Kwan *et al*. 2013). Natural variation in rDNA sequences and copy numbers are found even between closely related strains of yeast (James *et al*. 2009), yet almost nothing is known about how these differences influence any of the disparate functions attributed to the rDNA locus.

As our test system we chose to edit the rDNA origin of replication, an ~100 bp region between the divergently transcribed 35S and 5S genes in each repeat of the yeast rDNA array. One of the long-standing mysteries regarding the *rARS* is the observation that while each 9.1 kb repeat contains an identical *rARS* sequence, only a subset of repeats contains an active origin. The general inefficiency of the rARS on plasmids is thought to be due to the fact that the *rARS* ACS differs from the consensus sequence by a single G within one of the T tracts. An additional SNP is found in the ACS of the vineyard strain, RM11-1A, which reduces its plasmid maintenance properties even more. These non-canonical ACSs may help to explain the low efficiency of the *rARS* on plasmids; however, in the rDNA locus, transcriptional activity of adjacent rDNA repeats has also been shown to correlate with active origins (Muller *et al*. 2000). Furthermore, it isn’t known whether this origin inefficiency is a selected property, how it participates in or influences rDNA copy number, or how the status of the *rARS* influences the creation and/or segregation of extrachromosomal rDNA circles (ERCs) that have been associated with replicative aging. To begin addressing these questions, we have successfully replaced the rARS with both potentially more and less efficient origins and have characterized the key phenotypic effects of origin replacement.

To summarize our findings on Cas9 editing of the rDNA replication origin, we find that even with a single sgRNA, complete replacement of the rDNA ARS was efficient: when replacing the *rARS* with *ARS1* or *ARS1^max^*, we found that 8 of 12 colonies and 5 of 12 colonies tested (Figure S4B), respectively, had incorporated the introduced origin within the rDNA repeats. Deleting or introducing a loss-of-function mutation into the *rARS* appears less efficient (1-2 of 16 colonies; Figure S2C), perhaps due in part to viability problems that result from having no functional origin within the rDNA array. Regardless of which repair template we supplied, we found that cells respond to the severe reduction in rDNA copy number that results from Cas9 cleavage in one of three ways. 1) Transformants that fail to use a template for repair of the break delete the sequences between the most distal PAM sites and are unable to expand their rDNA locus beyond ~10 copies. Long-term growth selects for cells that have acquired an additional copy of chr II, chr IV or XII or have duplicated the entire rDNA locus through recombination at distantly located direct repeats (chr XII segmental aneusomy). 2) Transformants that repair their chr XII breaks with the *rARS^GC^* template can increase the number of tandem rDNA repeats and show a very modest return of origin activity to the NTS even though they retain only the GC mutant rARS. However, this level of expansion does not completely restore fitness as cells benefit from either whole chromosome disomy (chr II, IV or XII) or chr XII segmental aneusomy. 3) Transformants that have the “wt” rDNA template (or *rARS1* or *rARS1^max^*) for repair also experience a transient reduction in rDNA copy number. However, they are able to expand their rDNA locus, but within 100-150 generations do not reach the original ~150 repeats.

Based on our observations, using a variety of templates to repair Cas9 breaks in the rDNA, we suggest that the pathway of CRISPR/Cas9 editing is very similar to one proposed by Muscarella and Vogt (1993) for how cells survive cleavage of the rDNA locus by an endogenously expressed I-Ppo1 restriction enzyme. I-PpoI cuts once in each rDNA repeat and nowhere else in the *S. cerevisiae* genome (Lowery *et al*. 1992); however, surviving clones with mutations or insertions at the cleavage site are readily obtained. Upon sequencing, all of the repeats within a clone were found to share the same event, yet different clones experienced unique events. Muscarella and Vogt imagined a series of breaks that were repaired off a single mutant template, eventually sweeping through the 150 rDNA copies. While they did not examine rDNA copy number or chromosome XII sizes, it is likely that the I-PpoI survivors also experienced the reduction in rDNA size that that we have observed with Cas9 cleavage. In our experiments we provide repair templates that, once incorporated into the genome can serve as the template for repair of additional fragmented repeats.

Taking into account the evidence from the different rDNA editing experiments we performed we suggest the following model for how CRISPR/Cas9 editing leads to a homogeneous rDNA array (Figure 11): The ~20-fold difference in transformation frequency of pML104 and pACD suggests that Cas9 cleaves the PAM sites in the rDNA locus efficiently (Figure 11.1 and 11.2) and that for most cells cleavage is lethal. It is this lethality that allows us to detect repair without the need for a selectable marker. The rare cells that survived pACD transformation with their rDNA intact are likely to be the result of plasmid mutations that create a loss-of-function mutation in Cas9 or loss of the guides. For cells to survive cleavage of the rDNA repeats they would need to repair the broken chromosome by eliminating the target sites of the guide RNAs. We find that they repair the damaged locus in one of two ways: either the cut ends repair by nearby microhomology-directed recombination (Figure 12; Figure S3B) or they use the larger homology provided by the template with the altered PAM sites (Figure 11.3). The majority of the linear fragments (rDNA fragments and repair template) are lost through degradation (Figure 11.4). After rejoining the two halves of chr XII (Figure 11.5), it would only take the repair of a single excised repeat—by joining the two ends to form a circular DNA molecule—to create a more stable, non-cleavable template for repair of additional repeats, their reintegration into chr XII and expansion by unequal sister chromatid exchange (Figure 11.6). Our results with transformations using *rARS1* and *rARS1^max^* templates provide evidence for the rebuilding phase of the model: we found roughly a third of the clones had only ERCs and no appreciable *NTS2* sequences remaining on chr XII. The one clone we grew in liquid culture for an additional 170 generations demonstrated a rapid repopulation of the rDNA locus on chr XII (Figure S15B), presumably through integration of the ERCs. However, after rebuilding the chromosomal rDNA locus, these strains still maintain a much higher steady state level of ERCs—approximately 100-fold higher level than the wild type strain—while the strains with either *rARS^GC^ or rARS^Δ^* had no detectable ERCs. The lack of ERCs in strains with either *rARS^GC^ or rARS^Δ^* is likely the consequence of the inability of the ERCs to replicate once they are formed, although we cannot rule out the possibility that the status of the ARS is also playing a role in the ability of the ERCs to be excised from the rDNA locus.

**Figure 11:**
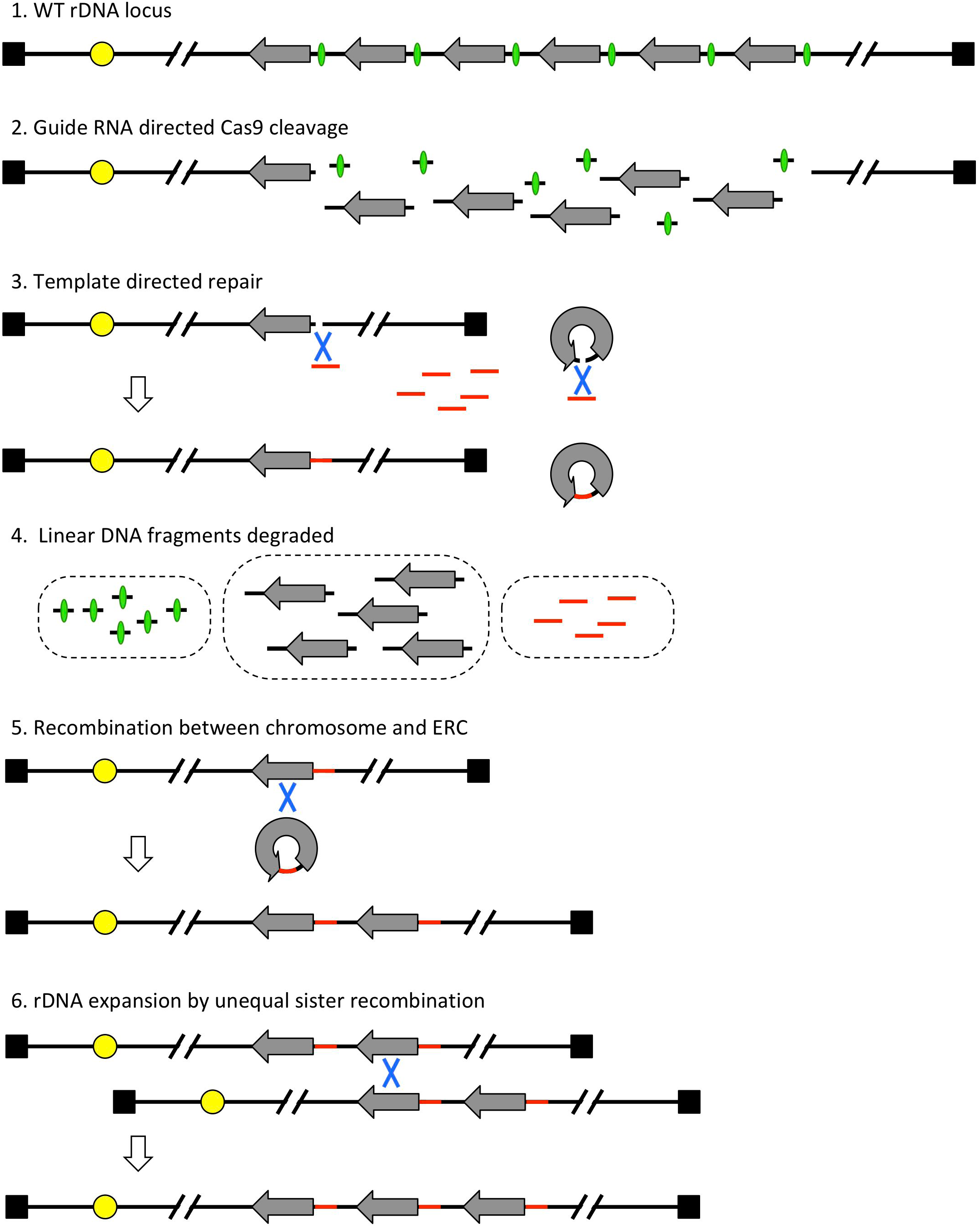
A proposed mechanism for CRISPR/Cas9 editing of the repeated rDNA locus. The excised rDNA NTS region that includes the origin is indicated by green ovals; the repair templates are indicated by red lines. The centromere of chr XII is a yellow circle and the telomeres are black squares.

**Figure 12:**
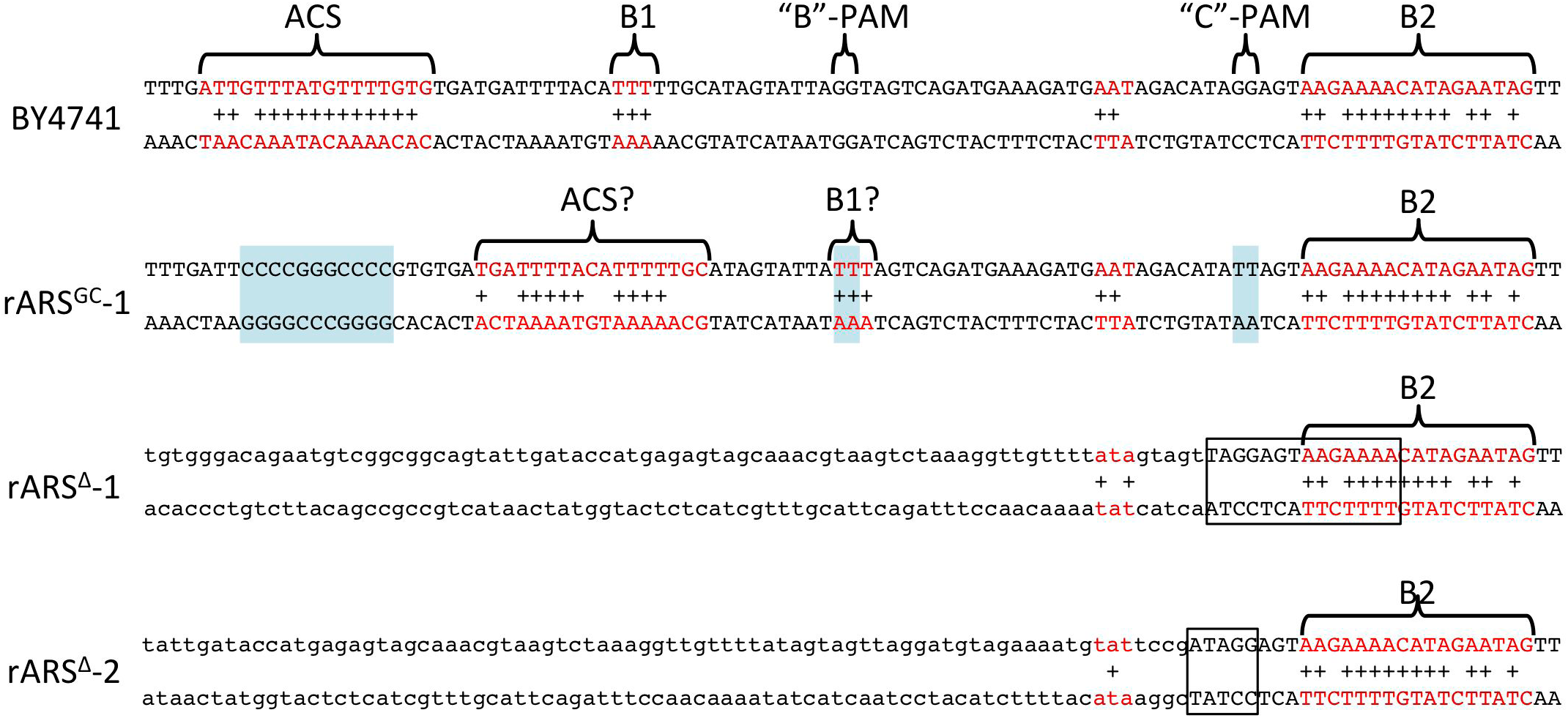
Identification of a cryptic origin in *rARS^GC^*. The core of the rDNA origin from BY4741 is shown with the ACS, the B1 elements and the B2 element in red type with a plus sign (+) indicating a match to the consensus sequences. (Note, this orientation of sequence is opposite to the orientation in Figure 1, to conform to the usual orientation of ARS sequences which places the T-rich strand of the ACS on the Watson strand to the left and the B2 element on the right.) Blue shaded boxes for rARS^GC^−1 highlight the GC replacement in *rARS^GC^* and the PAM site mutations associated with guides B and C. The brackets and red type highlight potential alternate ACS and B1 elements in *rARS^GC^*. rARS^Δ^−1 has a deletion of 167 bp and rARS^Δ^−2 has a deletion of 144 bp. Each deletion removes all four guide sequences and three of the four PAM sites. The lower-case nucleotides were originally 167 or 144 bp to the left of the junctions, respectively. The boxed sequences indicate the microhomology at the deletion junctions.

Clones that had been successfully edited came up slowly on transformation plates and retained their slow growth upon re-streaking on selective plates, suggesting that they may be compromised in their ability to synthesize rRNA. However, the only clones that maintain this slow growth on prolonged subculturing are those that have deleted *rARS* completely. It appears that the rDNA copy number that these strains are able to stitch back together reaches a maximum at ~10 copies. Limitations on replication are likely the cause of this ceiling. Since the rDNA locus is replicated unidirectionally, in the same direction as transcription, a single fork from the adjacent non-rDNA origin ~30 kb away would be responsible for replicating the entire 90 kb rDNA locus. Nowhere else in the yeast genome are potential origins of replication so distantly spaced (Raghuraman *et al*. 2001; http://cerevisiae.oridb.org). At an average fork speed of about 3 kb/min (Rivin and Fangman 1980), a 120 kb replicon would take longer than the ~30 minute S phase to complete replication. But there is reason to suspect that this fork would not be capable of the average speed because of the very high density of slower moving PolI transcription complexes (Schneider *et al*. 2007) that would be in the way (French *et al*. 2003; Ide *et al*. 2010). We envision two possible consequences of trying to replicate this rDNA locus from an adjacent origin: either the cell cycle would be delayed or cells would proceed into mitosis with an unfinished chr XII and suffer a double stranded break within the incompletely replicated rDNA locus. At ~45 generations we see significant (5%) chr XII breakage, which could reflect ongoing Cas9 editing or mitotic catastrophe from incomplete replication at the rDNA locus. The broken chr XIIs disappear upon continuous subculturing, suggesting that the cells have eventually completed the editing process or have solved the replication delay problem in some other way.

We have discovered several advantageous chromosomal amplifications that accompany long-term growth of the *rARS^Δ^* and *rARS^GC^* strains: aneuploidy for chr II, chr IV or chr XII and aneusomy for the region of chr XII that contains the rDNA locus. While we have not identified how chr II or chr IV aneuploidy improves fitness for the *rARS^Δ^* and *rARS^GC^* strains it is likely that some gene or genes on these chromosomes, when present in a second copy, could improve some aspect of DNA replication, protect cell cycle transitions or increase rRNA synthesis. The potential benefit of chr XII aneusomy/aneuploidy is more easily understood. By creating a direct repeat of the rDNA segment of chr XII between two directly oriented repeats or by maintaining a second copy of the whole chromosome, the cell doubles its ability to produce rRNAs without exceeding the ~120 kb ceiling on replicon size. Whatever detrimental effects result from having other genes in a second copy (Torres *et al*. 2007), they are countered by the benefit of increased rRNA production.

Unlike clones with *rARS^Δ^* that are unable to expand their rDNA copy number past ~10 copies, strains with *rARS^GC^* slowly increase rDNA copy number up to the proposed threshold required for efficient rRNA production (~35 copies; French *et al*. 2003; Kim *et al*. 2006). Simultaneously we found origin activity returning to the rDNA locus, but at a lower efficiency than in the parent strain, BY4741. We eliminated several possible explanations, including reversion to the wild type *rARS* sequence, creation of an origin sequence at a new site in *NTS2*, and trans-acting mutation elsewhere in the genome. However, closer examination of the sequence of *rARS^GC^* suggested to us that in the absence of the ACS, a new origin recognition complex (ORC) binding site (ACS + B1 elements; Figure 12) may have been created fortuitously by the repair template we used (in particular the “B”-PAM site mutation). Their absence in the *rARS^Δ^* sequences could explain why no origin activity is associated with *rARS^Δ^*. However, this hypothesis doesn’t explain why the new origin sequence in *rARS^GC^* clones remains inactive until the rDNA copy number expands beyond ~10. Two studies from the Diffley lab (Santocanale *et al*. 1999; Coster and Diffley 2017) may help explain the gradual return of origin function to the rDNA that we find on continuous culturing of *rARS^GC^*.

First, Coster et al. (2017) have refined our understanding of the role of the B2 element in origin function. They recently showed that the B2 element is likely to be a second ACS element and with their associated B1 elements are binding sites for two ORCs in inverted orientation. In this orientation and spaced ~70 bp apart they are able to recruit optimally the two head-to-head MCM complexes required for origin activation. When the distance between the two B1 elements is reduced to 50 bp, MCM loading in vitro drops by 50%, and to 30% when the distance is reduced to 25 bp. In the wild type *rARS*, the two B1 elements are 31 bp apart and for the proposed new ORC binding site, they are only 16 bp apart (Figure 12). If the results from their in vitro binding studies hold for the rDNA origin in vivo, then this spacing, combined with the less than optimal match to the ACS, may explain the low efficiency of the rDNA origin in the wild type sequence and its further reduction in the proposed new ACS/B1 sequence. Perhaps the closer spacing is what makes the wt *rARS* marginally functional and further reduces the substitute ACS/B1’s ability to establish the bidirectional MCM complexes.

Second, Santocanale et al. (1999) explored conditions that allow the dormant yeast origin *ARS301* to become active. They found that delay of forks from the adjacent *ARS305* is necessary to see *ARS301* become active, albeit weakly. Moreover, they showed that under these conditions, *ARS301* became active a full 45 minutes later than *ARS305*. Similarly, Vujcic et al. (1999) found that the normally inactive cluster of origins near the left end of chr III (including *ARS301*) became active in a strain that carried deletions of *ARS305* (25 kb away) and *ARS306* (59 kb away).

Putting together these two observations from the Diffley lab may explain the return of weak origin activity to the rDNA. The replacement of the original ACS with the G+C cassette and the introduction of the D-PAM site mutations have converted what remains of the *rARS* to a dormant origin. When there are ~10 repeats, all of the copies of the dormant *rARS* are passively replicated by the fork coming from the adjacent non-rDNA origin before they have a chance to become active; hence, there are no bubbles on 2D gels. However, during continuous culturing, the pressure to increase growth rate selects for cells that have increased their rDNA copy number. With 15 or 20 repeats, now one of the more distal *rARSs*, being further from the incoming fork, has time to fire before it is passively replicated; hence, low levels of rDNA bubbles become visible on 2D gels. This low level of initiation within the rDNA locus, along with the fork from the upstream, non-rDNA origin, could allow maintenance of the marginally expanded rDNA locus.

While we have explanations for some of the phenotypes associated with editing the *rARS*, there are still some interesting questions that remain for future studies. First, *rARS1* and *rARS1^max^* are both much more efficient plasmid and chromosomal origins than the wild type *rARS*, yet, when inserted into the rDNA *NTS2*, they are less efficient in that environment than the *rARS* (Figure 2). What features of chromatin structure, nucleolar location or transcriptional activity are influencing these origins differently than the native *rARS*? Second, wild type BY4741 normally has about 150 copies of the rDNA, yet even replacing the *rARS* with *rARS^“wt”^*, the copy number is very slow to expand and doesn’t begin to reach this large number of repeats even after 150 generations of continuous selection. This result contrasts with the more rapid expansion observed by Kobayashi et al. (1998) upon restoration of functional Rpa135 to a W303-derived strain in which the rDNA locus had been shrunk due to loss of Rpa135. Third, how does the status of the *rARS* contribute to fitness and lifespan? The potential efficiency of the origin in the rDNA is clearly correlated with the ability of the cells to create and maintain ERCs, with more than a 100-fold difference between *rARS^GC^* or *rARS^Δ^* and *rARS1* or *rARS1^max^*, yet all of the edited strains have severely shortened lifespans, regardless of ERC level. Perhaps the drastic reduction in rDNA copy number produced by CRISPR/Cas9 editing results in a ribosome deficiency that masks any effects that ERCs or origin efficiency might have on lifespan. Alternatively, having so few templates for transcription could be creating a burden that interferes with replication or repair of the rDNA, and perhaps it is one of these processes that is responsible for the reduction in lifespan. Discovering the specific gene or genes on chr II or IV that increase fitness and/or lifespan when duplicated will help to resolve these questions. Finally, single-copy origins elsewhere in the yeast genome are individually dispensable. Even the deletion of adjacent origins on chr III (Dershowitz *et al*. 2007) and chr VI (Bogenschutz *et al*. 2014) causes no obvious growth defects. These findings have led to the conclusion that “no single origin is essential”. Yet we find that deletion of the rDNA origin has profound consequences on fitness and lifespan by restricting the number of rDNA repeats that can be passively replicated from the adjacent upstream origin and thus reducing ribosome production. So, is the rDNA origin the exception to the rule that “no single origin is essential”? And if the rDNA origin is so critical to rDNA maintenance, then why is the origin inherently so inefficient? What are the evolutionary constraints on ORC and the rARS that drive this sub-optimal interaction?

We have shown that CRISPR/Cas9 editing of a tandem repeated locus can be efficient and informative, but we have only edited the origins. Many other interesting functions have been mapped to the rDNA repeats of yeast, both to transcribed and non-transcribed regions. For example, the RFB that enforces the unidirectional traffic of RNA and DNA polymerases in the rDNA has been studied largely by analyzing mutations in Fob1, the protein that binds the RFB and carries out various functions associated with rDNA copy number expansion, recombination, ERC production and lifespan determination (Kobayashi *et al*. 1998; Kaeberlein *et al*. 1999; Takeuchi *et al*. 2003). CRISPR/Cas9 technology would make the mutational analysis of the RFB in situ a fruitful parallel avenue of study. There are also PolII promoters and transcription units in the rDNA repeats that are well documented and positioned in interesting places where they may be influencing replication, recombination, sister chromatid cohesion or PolI transcription (Poole *et al*. 2012; Saka *et al*. 2013). It should now be possible to edit these promoters to assess more completely their roles in rDNA metabolism. Transcriptional silencing by Sir2 and loading of cohesion/condensin in the rDNA play important roles in many aspects of rDNA metabolism and cell health (Kaeberlein *et al*. 1999; Johzuka *et al*. 2006; Fine *et al*. 2019) that have yet to be fully explored through alterations to the DNA template. In addition, using CRISPR/Cas9 technology it should now be possible to explore the role of unstructured loops and less-well conserved regions in the rRNAs on ribosome function in vivo (for example, Fujii *et al*. 2018). Finally, the technology could be extended to higher eukaryotes to genetically dissect their highly repetitive loci—including rDNA, centromeres, segmental duplications and other tandemly repeated loci yet to be discovered.

## Supplemental Figure Legends

**Figure S1:** Maps of CRISPR/Cas9 plasmids and PCR fragments from the *NTS2* region of wild type and edited strains. A) Plasmid pML104 was modified by the insertion of three tandem sgRNA templates (grey arrows marked A, C, and D) or a single sgRNA template (F). B) The 762 bp wild type rARS region amplified by PCR primers (arrows) shows the relevant locations of PAM sites (A, B, C, D, and F), the ACS (green box) and a unique HaeIII site. Sizes (in bp) refer to the HaeIII cleavage products. Each of the Cas9-edited *NTS2* regions produces unique PCR fragment sizes and/or restriction fragment sizes. The ~100 bp insertions of *ARS1* or *ARS1^max^* are indicated by the inverted triangles.

**Figure S2:** Isolation and confirmation of Cas9 edited rDNA repeats. A) BY4741 was transformed with pML104 (control plasmid with only Cas9) or pACD (three sgRNA + Cas9) and the *rARS^GC^* repair template in either single stranded or double stranded form. B) Colony purification of eight transformants from the plates shown in A. Clones maintain their fast/slow growth properties. C) Characterization of PCR products of the *NTS2* region. Replacement of the rDNA origin with *rARS^GC^* produced the same size fragment as the wild type sequence but was sensitive to XmaI cleavage. The symbols below the XmaI cleavage gels summarize the events: **+** = un-edited sequence; Δ = deletion of all or part of the fragment targeted by PCR; ✔ = correct replacement;= incomplete digest or mixed PCR products. The clones rARS^Δ^−1 and rARS^GC^−1 are indicated with arrows.

**Figure S3:** Sanger sequence trace of rARS^GC^−1 (A) and rARS^Δ^−1 (B). A) The sequence of the Watson strand from BY4741 (Chr XII coordinates in bp), taken from the *Saccharomyces* Genome Database, is shown at the top with the ACS element and the A PAM site indicated with brackets. The sequence trace the *rARS^GC^* shows the successful replacement of both sequences. A blow-up of the region encompassing the GC-replacement illustrates no evidence of the original ACS remaining. B) The sequence of the Watson strand from BY4741 aligned to highlight the region of homology involved in the rARS deletion (gray box). The sequence trace confirms the breakpoint in the deletion and reveals that the rDNA repeats in rARS^Δ^−1 are a mixture of two sequences differing by 3 bp in their junctions.

**Figure S4:** Confirmation of *rARS* editing by PCR and restriction digestion. A) BY4741 was transformed with pACD and *rARS^”wt”^* repair template that has all four PAM site mutations. PCR fragment sizes reveal six potential deletion strains. Restriction digestion of the PCR products distinguishes unedited clones (+) from edited clones (✔) by differential digestion at the locations of the two PAM sites A and D. ApoI cleaves the edited A PAM site and HaeIII cleaves the unedited PAM site D. B) rARS^GC^−1 was transformed with pF and *rARS1^max^* repair template. PCR and restriction digestion confirms that six of eight transformants were correctly edited (✔) while one retained the original GC insert (**+**) and one produced a mixed digestion pattern (**?**). C) BY4741 was transformed with pACD and either *rARS1* or *rARS1^max^* repair template. PCR and restriction digestion with BglII and BsaA1 distinguish the clones replaced by *rARS1* or *rARS1^max^*, respectively. In addition to unedited clones (+), correct transformants (✔) and apparent mixed clones (?), we recovered a deletion clone (Δ). Clones 1 and 2 of the *rARS1* transformants and clones 1-4 of the *rARS1^max^* transformants produced large colonies on the transformation plates. For future analysis of transformants, we only characterized small colonies.

**Figure S5:** CHEF gel analysis of BY4741 clones transformed with pML104. A) Ethidium bromide stain of the CHEF gel containing eight transformants shows little variation in chr XII size. B) and C) Southern blots of the CHEF gel in A probed sequentially with *CEN12* and *MAS1* show there is no prominent sub-chromosomal sized fragments from chr XII. The faint patterns below chr XII are cross hybridization to the other yeast chromosomes.

**Figure S6:** Chromosomal location of rDNA sequences in Cas9 transformants. CHEF gel Southern blots from Supplemental Figures 2 and 5 were reprobed with an *NTS2* probe to look for translocation of rDNA sequences to other chromosomes or as extrachromosomal molecules.

**Figure S7:** CHEF gel analysis of *rARS^”wt”^* transformants. A) The ethidium bromide photograph is shown along with Southern blots using *MAS1* and 37S probes to assess chr XII size and rDNA repeat locations, respectively. Each transformant is categorized based on PCR and restriction digests shown in Supplemental Figure 4C. B) Genomic DNA from the same 12 transformants was cleaved with FspI in agarose plugs. The rDNA copy number of each strain was deduced from the FspI fragment sizes on the Southern blot of the CHEF gel using uncut yeast chromosomes as size markers. C) FspI digests of *rARS^Δ^* and *rARS^GC^* haploids and heterozygous diploids.

**Figure S8:** Estimating 25S rRNA content of rARS^GC^−1 by comparative hybridization of a single copy gene, *ACT1*. A) Total nucleic acids were isolated from log phase cultures of BY4741 and rARS^GC^−1 and separated on an agarose gel. Lanes 2 and 4 contain nucleic acids from BY4741; lanes 3 and 5 contain nucleic acids from rARS^GC^−1. B) The gel was cut into two portions. The upper portion was Southern blotted and the lower portion was blotted as a northern. The Southern blot was probed for *ACT1* and the northern blot was probed for 25S rRNA. C) Quantification of total counts for replicate lanes were tallied and a ratio of rRNA/ACT1 for rARS^GC^−1 was normalized to BY4741.

**Figure S9:** Increase in chr XII size over 100 generations in three isolates with *rARS^”wt”^* (A) and one isolate of *rARS1^max^* (B).

**Figure S10:** Selection for suppressors of slow growth in turbidostat cuptures of one *rARS^Δ^* and three *rARS^GC^* clones. A) The growth rates calculated from the pump speeds for strains over the last ~five days of the turbidostat runs. B) CHEF gels for samples harvested on day 3 and on the days indicated in (A) by the black triangles. C) Southern blot analysis of changes in chr XII/rDNA over the course of the continuous growth.

**Figure S11:** Detecting chr II aneuploidy by CHEF gels. A) and B) The CHEF gels in Figure 4C were simultaneously hybridized with probes near CENs 2 and 9. C) Quantification of the ratio of chr II to chr IX hybridization signals were normalized to that of BY4741. Samples from the same DNA plugs for BY4741 and rARS^GC^−1 were included on both gels.

**Figure S12:** Monitoring chr XII size during serial passage of rARS^GC^−2 and rARS^Δ^−2. A) The ethidium bromide stained CHEF gel contains samples from the serial passage and four independent clones isolated at generation 90. B) Samples from alternate days of the same serial passage experiment were run under conditions to reveal subtle changes in rDNA sizes. The Southern blots were probed sequentially with *CDC45* to detect chr XII and then with *CEN2* and *CEN9*, simultaneously, to assess chr II copy number. C) and D) The Southern blot of the gel from A was probed sequentially with *CDC45* to detect chr XII and then with *CEN2* and *CEN9*, simultaneously, to assess chr II copy number. E) Quantification of the ratio of chr II/chr IX over the course of the continuous growth and in the gen-90 clones from a CHEF gel with samples collected every 10 generations.

**Figure S13:** Read-depth analysis from WGS for evolved rARS^Δ^−2 and rARS^GC^−1 clones. Read depth from WGS of genomic DNAs from populations on the last day of Run1 and Run2 turbidostat populations (T1-T4) reveals an increased copy number for chr IV or chr XII or a partial duplication of chr XII that includes the rDNA locus. Read-depth ratios of Chr IV or Chr XII were compared to ratios across ChrV, VI, VII and IX using the Wilcoxon-Rank Sum test. Ratios with a P value of >0.0001 are indicated by their relative copy number differences above each significantly amplified chromosome or chromosomal segment.

**Figure S14:** Analysis of clones transformed with pF and *rARS1* or *rARS1^max^*. A) CHEF gel analysis and Southern blotting of eight clones transformed with *rARS1^max^*. Notice the different hybridization patterns for the *MAS1* and 37S probes for clones 2, 4, and 7. B) CHEF gel analysis and Southern blotting of eight clones transformed with *rARS1*. Notice the different hybridization patterns for the *MAS1* and 37S probes for clones 2 and 5. C) 2D gel analysis of rARS1^max^−2 and rARS1^max^−4 that contain predominantly circular forms of the rDNA. The red circle highlights the absence of the 22.4 kb junction fragment between the rDNA locus and unique chr XII sequences.

**Figure S15:** CHEF gel analysis of Cas9 edited clones with circular rDNA. A) rARS1^max^−4 was restreaked and 12 individual colonies were analyzed to determine the stability of the extrachromosomal circular rDNA. The CHEF gel and Southern blot indicate that the ERSc are stably inherited. B) rARS1^max^−4 was serially passaged in liquid culture for 170 generations and analyzed by CHEF gel and Southern blotting. Samples at 10, 30, 50, etc. generations were examined. By 30 generations cells with stably integrated rDNA repeats had overtaken the culture.

## Supplemental Tables

**Table S1: List of Primers, oligonucleotides, gBlocks and Sanger sequencing**

**Table S2: List of variant calls from WGS**

**Table S3. Variant filter parameters**

**Table S4. Summary of chromosomal/rDNA amplification events**

